# The Genetics of Participation: Method and Analysis

**DOI:** 10.1101/2022.02.11.480067

**Authors:** Stefania Benonisdottir, Augustine Kong

## Abstract

Participation in a genetic study likely has a genetic component. Identifying such component is difficult as we cannot compare genetic information of participants with non-participants directly, the latter being unavailable. Here, we show that alleles that are more common in participants than non-participants would be further enriched in genetic segments shared by two related participants. Genome-wide analysis was performed by comparing allele frequencies in shared and not-shared genetic segments of first-degree relative pairs of the UK Biobank. A polygenic score constructed from that analysis, in non-overlapping samples, is associated with educational attainment (*P* = 2.1 × 10^−52^), body mass index (*P* = 1.5 × 10^−19^), and participation in a dietary study (*P* = 6.9 × 10^−21^). Further analysis shows that inclination to participate is a behavioural trait in its own right, and not simply a consequence of other established phenotypes. Understanding the basis of this trait is important for data analyses and the design of future surveys, genetic or otherwise.

## Introduction

For all sample surveys, ascertainment bias, *i.e*. the sample is not representative of the population, is a problem that could lead to seriously misleading conclusions^1,2^. By its very nature, ascertainment bias usually cannot be evaluated based on the sample alone^3^. For example, with a target variable such as the presidential candidate one is voting for or whether one has been vaccinated, other variables (covariates) that have known distributions for both sample and population are needed for potential adjustments^1–3^. Such adjustments are inherently imperfect as the covariates are unlikely to fully capture the correlation between participation and the target variable^1,3^. For genetic investigations, among participants of the primary study who have contributed DNA, further engagement in optional components of the study has been demonstrated to have associations with both genotypes and phenotypes^4–7^. That, however, does not address the potential genotypic difference between the primary study participants and the target population. Given this background, it is striking to see that it is actually possible to investigate how the sampled genotypes are biased based on themselves alone. A recent study identified single nucleotide polymorphisms (SNPs) that had significant allele frequency differences between males and females in the samples, and proposed that those variants have differential participation effects for the sexes^8^. This relies on the assumption that genotypes of autosomal variants should have the same frequencies in the population for both sexes. This approach, however, cannot identify variants that affect primary study participation of both sexes in a similar manner, which are presumably more abundant and have broader impact. Here we show a way to study these variants.

There are two properties that make genetic data distinct. Firstly, all individuals are genetically related by various degrees, and the relatedness are captured by the genotypes. Secondly, each individual has two copies of genetic segments on autosomal chromosomes, one inherited from each parent. Some of these segments are identical by descent (IBD), *i.e*. inherited from a recent common ancestor, with genetic segments in a relative. Conceptually, instead of comparing individuals, we compare genetic segments. The key idea introduced here is that if an allele has a higher frequency in participants than non-participants, it would also have higher frequency in segments that are in two participants compared to segments that are only in one participant. This alternative view of the data leads to the three principles of genetic induced participation bias described below. These principles allow us to perform a genome-wide association scan (GWAS) of study participation variants using only the genetic data of the study without phenotypes. Most importantly, this analysis is only sensitive to direct genetic effects^9–11^, and is immune to indirect genetic effects and confounding effects such as those induced by population stratification^12^. Examination of the GWAS results reveals that, while a person’s participation is related to other characteristics such as educational attainment, the effect of the genetic component of participation is not simply manifested through them. This partially explains why standard covariate adjustment cannot fully eliminate the effect of ascertainment bias in general.

### The First Principle of Genetic Induced Ascertainment Bias

> On average, between two ascertained individuals, genetic segments shared identical by descent (IBD), relative to segments that are not, are enriched with alleles that have positive direct effects on ascertainment probability.

This principle (Fig. 1A) relies on individuals in the population being genetically related, closely or distantly. Gene alleles that are enriched in the participants compared to the non-participants are expected to be further enriched in segments shared by ascertained relatives. With a large sample, at a specific SNP locus, many pairs of individuals would share one long haplotype, inherited identical by descent from a not-very-distant common ancestor. For each of such pairs, there is one distinct shared haplotype, and two distinct not-shared haplotypes. Tabulating over all such pairs the SNP alleles in the shared haplotypes and the not-shared haplotypes, the SNP allele that promotes participation/ascertainment should tend to have a higher frequency in the shared than the not-shared haplotypes. This becomes a case-control analysis where the shared and not-shared alleles are the cases and controls respectively, and matched to the extent that they are in the same individuals. Still, that does not remove potential confounding entirely as haplotypes in the population that were driven to higher frequency through natural selection would also be shared by more individuals, both in the population and in the sample. Essentially, ascertainment bias is a form of selection and to cleanly distinguish it from other forms of selection requires more stringent matching of shared and not-shared haplotypes. We achieve that by using ascertained parent-offspring and sibling pairs.

**Figure 1.**
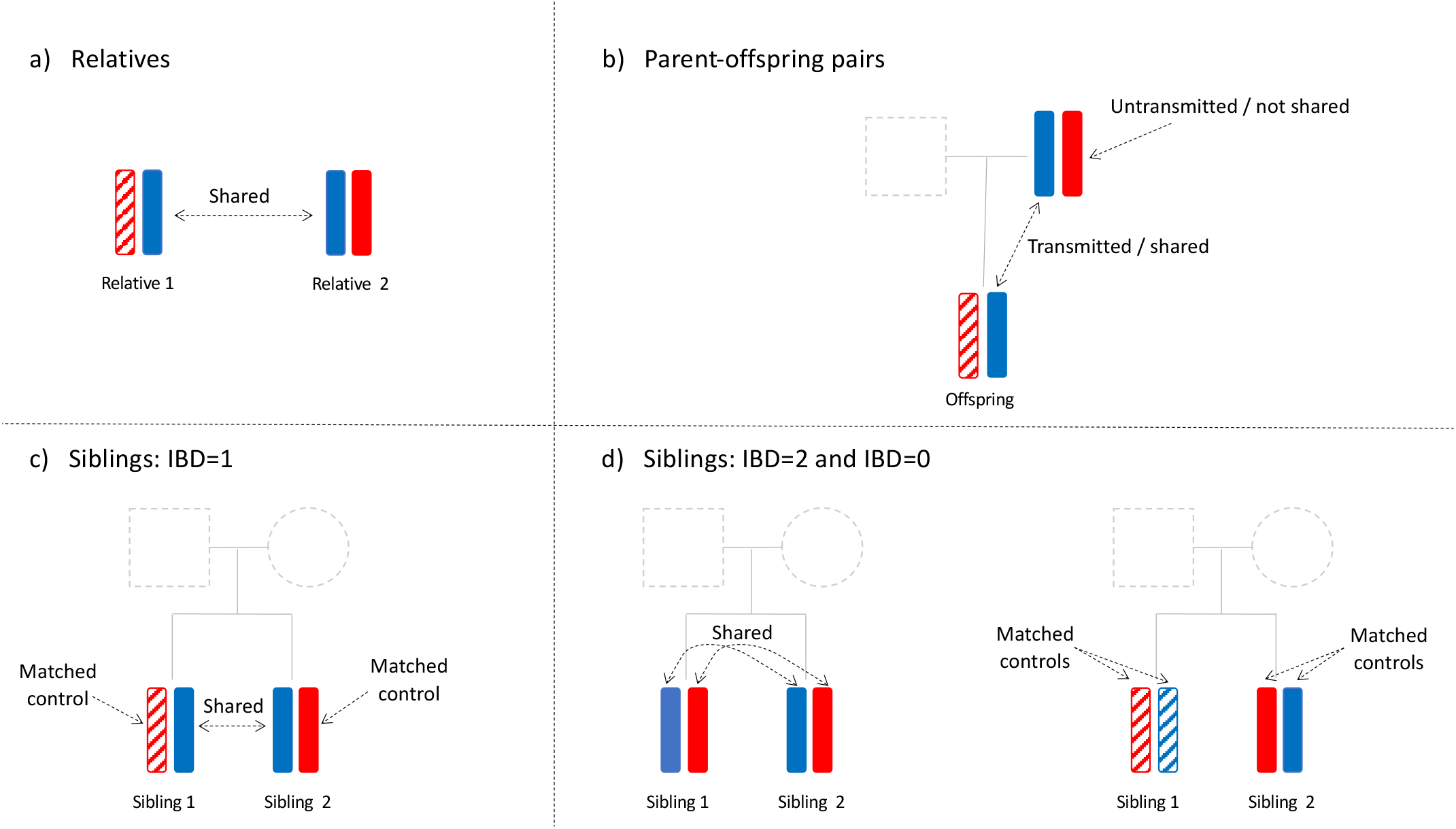
The First Principle for Genetically Induced Participation Bias; comparing shared and not-shared alleles. a) General relative pairs. b) Parent-offspring pairs. c) Sibling pairs with IBD = 1 at SNP locus. d) Sibling pairs with IBD = 2 and IBD = 0 at SNP locus.

### Ascertained parent-offspring pairs (Fig. 1B)

For an ascertained parent-offspring pair where the other parent is not ascertained, there are three distinct alleles by descent, (*S*) the allele in the genomic segment transmitted from parent to offspring, (*NS_P_*) the allele in the parental genomic segment not transmitted to the offspring, and (*NS_OP_*) the allele inherited by the offspring from the other parent. Thinking of the offspring as the proband, the *NS_P_* allele is a perfect match for the *S* allele as they are both in the ascertained parent. Mendelian inheritance dictates that each would have the same chance to be transmitted to the offspring to become the shared allele. The principle is similar to that underlying the transmission disequilibrium test^13^. The *NS_OP_* allele is not as perfectly matched, and while it could be used for subsequent validation, it is not used in our initial analysis. With alleles coded as 0/1 and *n_PO_* ascertained parent-offspring pairs,

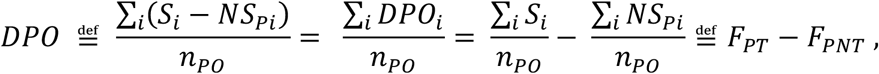

where *i* = 1,…, *n_PO_* indexes the pairs, can be used to test for association between SNP and ascertainment. In particular, conditional on the genotypes of the ascertained parents, *DPO* has expectation zero under the null hypothesis of no ascertainment bias, and is not subject to any population stratification induced bias. The case where both parents are ascertained together with an offspring can be treated as two parent-offspring pairs as the transmission from the two parents are independent under the null hypothesis.

### Ascertained sib-pairs, parents not ascertained

Let *n_SIB_* be the number of ascertained sib-pairs with unascertained parents. At a specific locus, assuming random Mendelian transmission, a sib-pair has probability ½ of inheriting the same allele IBD from the father, and the same independently from the mother. It follows that a sib-pair would share 2, 1, or 0 alleles IBD with probabilities ¼, ½ and ¼ respectively. With data from dense SNPs, the IBD state of a locus can usually be determined with high accuracy^14^. For a specific SNP, based on the sib-pairs with IBD states that are assumed known, two frequency-difference statistics are derived as follows. For each sib-pair that has IBD state 1 (Fig. 1C), there are one distinct share allele (*S*) and two distinct not-shared alleles (*NS*_1_ and *NS*_2_). Define

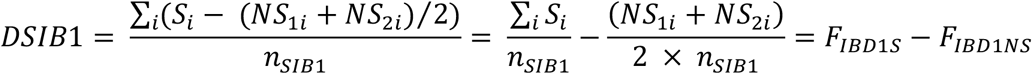

where *i* =1,…, *n_SIB1_*. Note that for any one of these sib-pairs, if the shared allele is paternally inherited, then the two not-shared alleles are maternally inherited, and vice versa. Despite that, conditional on the genotypes of the two parents, without ascertainment bias, *DSIB1* has expectation 0. This holds even in the most extreme case where fathers carry only allele 1 and mothers carry only allele 0. That can be demonstrated by considering the four parent-offspring transmissions --- one paternal transmission for each sib and one maternal transmission for each sib --- jointly (Fig. S1). Notably, when the IBD state is 1, the shared allele is equally likely to be paternal or maternal, and thus any systematic differences between fathers and mothers cancel in expectation. For the approximately one-quarter of the sib-pairs (*n_IBD2_*) at a locus that share two alleles IBD, the average allele frequency is

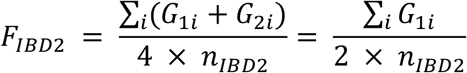

where *i* = 1,…, *n_SIB2_*, indexes the sib-pairs with IBD state 2, and *G_1i_* and the identical *G_2i_* are the genotypes of sib 1 and sib 2 respectively in pair *i*. Similarly, for the approximately one-quarter of the sib-pairs (*n_IBD0_*) at a locus that share zero alleles IBD, let

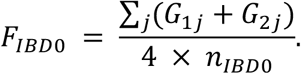

where *j* = 1,…, *n_SIB0_*, indexes the sib-pairs with IBD state 0, and *G_1j_* and *G_2j_* are the genotypes of sib 1 and sib 2 respectively in pair *j*. The difference (Fig. 1D)

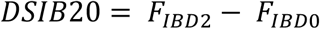

is another test statistic for ascertainment bias. Here *F_IBD2_* and *F_IBD0_* are allele frequencies from different sib-pairs. However, for a sib-pair at a particular locus, the chance to be in IBD state 0 or 2 is the same, and thus, without ascertainment bias, *DSIB20*, has expectation zero.

In summary, *DPO*, *DSIB1*, and *DSIB20* are only sensitive to the direct effects^9,15^ on ascertainment, and are unaffected by population stratification induced bias^12,15^ or indirect genetic effects from relatives ^9–11^. However, direct genetic effects can manifest in various ways, their relative contributions depending on the sampling scheme. For example, the effect can be ‘voluntary’, a person is more likely to participate when invited, or ‘involuntary’, *e.g*. a person is less likely to be invited because of its attributes. The three statistics can be combined, *e.g*. as a weighted linear combination, into one test statistic. However, having separate values of them are useful because they can have different expectations depending on the nature of the ascertainment bias, and they are impacted differently by genotyping and data processing errors.

## UK Biobank and Data Processing Artefacts

The UK Biobank (UKBB) is a large-scale database with genetic and phenotypic information of individuals from across the United Kingdom (UK)^16^. Invitations to participate were sent to 9,238,453 individuals who were aged between 40 and 69 years and lived within 25-mile radius of any of the 22 UKBB assessment centres^17^. In the end, 5.45% of those participated in the study (~500,000 individuals) and went through baseline assessments that took place from 2006 to 2010.^17^ In addition to phenotypic details collected at the baseline visit, information continued to be added, including follow-up studies for large subsets of the cohort^17–19^. It is known that the UKBB sample is not fully representative of the UK population^16,17,20^. The participants were more likely to be female^17^, less likely to smoke^17^ and were older^17^ than non-participants. Also, compared to the national average, participants were more educated^20^, taller^17^ and had a lower BMI^17^.

We applied our methods to 4,427 parent-offspring pairs and 16,668 sibling pairs of white British (WB) descent (Fig. S2, Table S1 and Supp. Text). Association analysis was performed for 597,039 directly genotyped phased SNPs^16^ that passed quality control filtering (Supp. Text) and had a minor allele frequency (MAF) > 1%. For each sibling pair, IBD sharing status (0, 1 or 2) of every SNP was ‘called’ using the program KING^14^ (Supp. Text). Initial results had the major allele of a SNP significantly more likely to be positively associated with participation. Examination of possible artefacts exposed a few data processing issues, leading to adjustments outlined below.

For many loci, the IBD fractions of the sibling pairs, calculated from the calls, deviated significantly from the expected IBD fractions (Supp. Text, Fig. S3) indicating IBD calling errors. Errors were most likely to occur around recombination events and ends of chromosomes (Fig. S3) and to reduce their impact, for each sibling pair, we identified the IBD segments --- contiguous SNPs with the same IBD status called --- and trimmed away 250 SNPs from each end of a segment before the association analysis. This removed 250 SNPs at each end of the chromosomes entirely from the association analysis and reduced slightly the sample size of the other SNPs (Fig. S4, median reduction of 1,516 pairs).

For a parent-offspring pair and for SNPs that have IBD status 1 for a sibling pair, the shared allele and not-shared alleles are clear unless both members of the pair are heterozygotes, in which case the phasing with neighbouring SNPs was used to resolve the uncertainty (Fig. S5 and Supp. Text). Our analysis estimated that the phasing provided in the UK Biobank data release^16^ had an error rate of approximately 0.5% and an adjustment was made to the test statistics *DPO* and *DSIB1* to account for the induced bias (Supp. Text).

After the above adjustments, the tendency for the major allele to be positively associated with participation was substantially reduced but not completely eliminated. Even though part of this ‘major allele effect’ might be real, we made further adjustments based on MAF to remove this tendency from our association results (Supp. Text). After all these adjustments, a number of highly significant associations remained in the major histocompatibility complex (MHC), a region with extended linkage disequilibrium (LD) and high SNP density. Further investigations led us to believe these signals were most likely artefacts (Supp. Text and Fig. S6). After removing 152,582 SNPs from the MHC and other extended LD regions (Table S2) as well as additional quality control filtering (Supp. Text and Fig. S7), 444,457 SNPs remained in our participation genome scan.

## Results

For each SNP, we computed three t-statistics by dividing each of *DPO, DSIB1*, and *DSIB20*, by its standard error (SE). The three t-statistics were then combined into one, based on their SEs, and then converted to a nominal chi-square statistic. The latter were adjusted by genomic control using the genomic inflation factor. A QQ-plot of the p-values (*P*) based on the adjusted chi-square statistics (Fig. S8) shows that one SNP, rs113001936 on chromosome 16, is significant with Bonferroni correction, and the excess of SNPs with *P* < 0.01. A literature search did not provide any obvious reasons for rs113001936 to have a participation effect. To validate our results, we derived the weights of a participation polygenic scores (*pPGS*) based on our participation GWAS (Supp. Text). Values of the *pPGS*, standardized to have variance one, were computed for 272,409 WB individuals with no close relatives in UKBB (>3^rd^ degree for all pairs). They are referred to as the ‘unrelateds’ and notably do not overlap with the first-degree relatives used. Associations between the *pPGS* and various quantitative traits were examined using the unrelateds. Table 1 shows some of the strongest associations and includes a few non-significant ones. The strongest association is with EA where the effect (correlation) is 0.0307 with *P* = 2.1 × 10^-52^. The effect, 0.0299, is nearly as strong for age-at-first-birth (AFB) of women (*P* = 1.2 × 10^-20^). The next strongest association is with BMI (*P* = 1.5 × 10^-19^) where the effect is notably negative. These results, consistent with the known differences in EA and BMI between sample and population^17,20^, validated that our GWAS performed without any phenotype data can nonetheless capture genetic associations with participation.

**Table 1:**
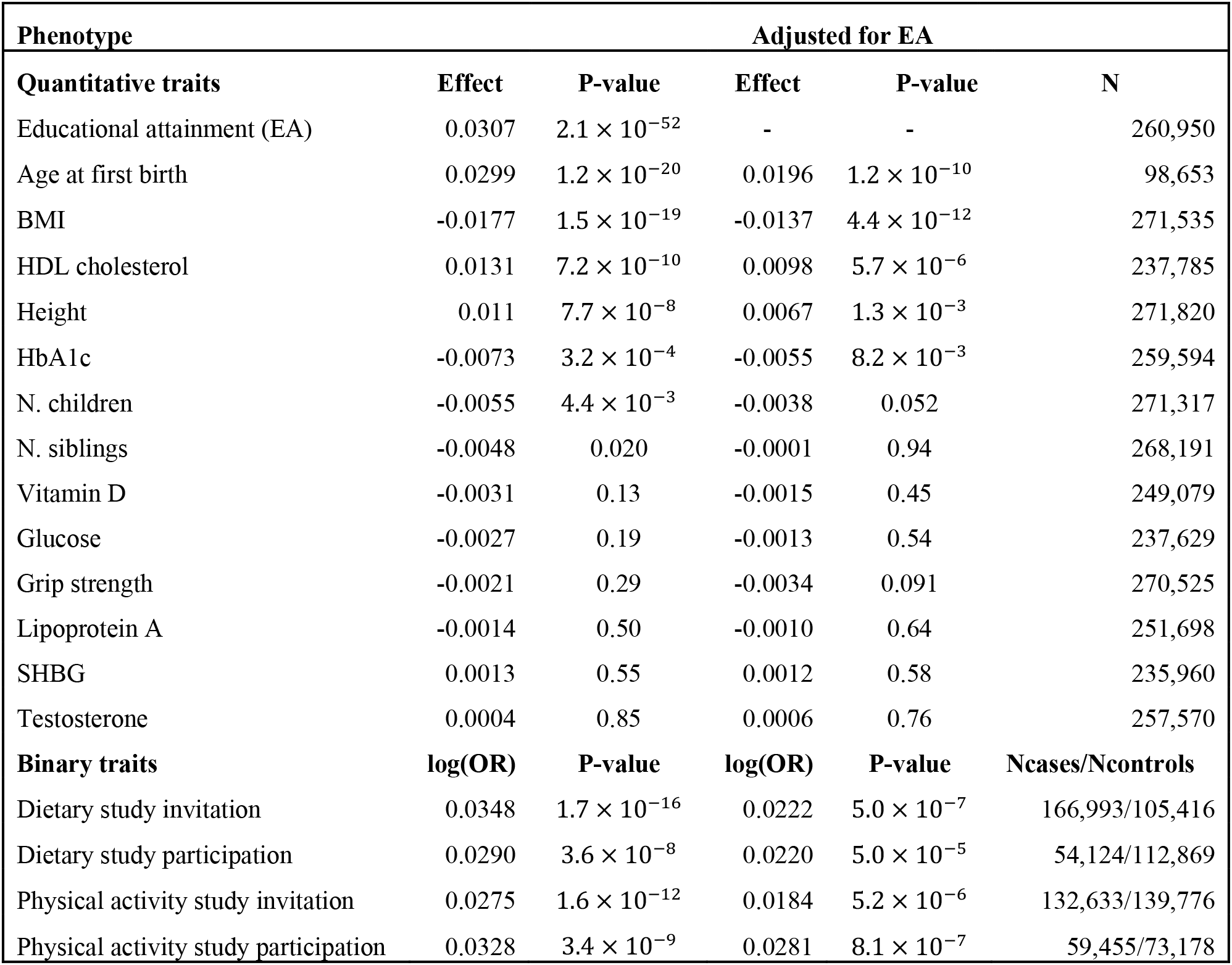
Participation PGS and association with phenotypes. This table depicts how the participation polygenic score (PGS) is associated with a range of phenotypes. For the quantitative traits, we performed linear regression with the corresponding phenotype as a response and the PGS and genotyping array (BiLEVE or Axiom) as explanatory variables in the subset of individuals of White British descent who have no close relatives in UK Biobank (>3^rd^ degree for all pairs). Prior to regression analysis, the quantitative traits were adjusted for year of birth (YOB), age at measure and 40 PCs separately for each sex and then the residuals were standardised also separately for each sex (Supp. Text). Grip strength was additionally adjusted for height. BMI: body max index. HDL cholesterol: High-density lipoprotein cholesterol. HbA1c: Glycated Haemoglobin. N. children: Number of children. N. siblings: Number of full siblings. SHBG: Sex hormone binding globulin. Information about age at first birth was only available for women. For the binary traits, we performed a logistic regression including YOB, age at measure up to the order of three, 40 PCs, sex and genotyping array as additional covariates. Columns 2 and 3, labelled ‘Effect/log(OR)’ and ‘P-value’, depict the slope/log(odds-ratio) for the participation PGS and the corresponding P-value. The P-values have been adjusted with the LD-score regression intercept of the corresponding phenotype (Supp. Text). Columns 4 and 5 show the slope/log(odds-ratio) and the P-value for the participation PGS when educational attainment (EA) has been added as an additional covariate. SNP-wise weights for the participation PGS were computed using identity-by-descent information from 16,668 siblings and 4,427 parent-offspring pairs in UK Biobank.

After the baseline assessments, subsets of the UKBB participants were invited to further participate in a dietary study in 2011-2012 where invited participants were asked to answer a dietary questionnaire^19^ and a physical activity study in 2013-2015 where invited participants were asked to wear an accelerometer for a week^18^. Not everybody was invited (the criteria for invitation included having a valid email address) and only a subset of those invited actually participated (Table S3). We refer to participation in these follow-up studies as ‘secondary participation’^4–7^. For the dietary study, the estimated effect of *pPGS* on being invited, in loge odds-ratio (*log*(*OR*)), is of 0.0348 (*P* = 1.7 × 10^-16^, Table 1). For those invited, the estimated effect on actual participation in log(*OR*) is 0.0290 (*P* = 3.6 × 10^-8^). For the physical activity study, the corresponding effect estimates are 0.0275 (*P* = 1.6 × 10^-12^) and 0.0328 (*P* = 3.4 × 10^-9^) respectively. Comparing the *pPGS* values of the three groups --- uninvited, invited but did not participate, participated ---, a 2-*df* test gives *P* of 6.9 × 10^-21^ and 8.5 × 10^-19^ respectively for the dietary and physical activity studies.

Associations of *pPGS* were further examined with adjustment for EA (Table 1). For traits/variables with unadjusted *P* < 1 × 10^−3^, the adjusted effects shrink but all remain significant. Relatively, the effect on participating in the physical study when invited shrinks the least, by 14.3%, from 0.0328 to 0.0281, while the effect on dietary study invitation shrinks by 36.2%, from 0.0348 to 0.0222. This difference in shrinkage is partly because the estimated effect of EA on dietary study invitation is 0.4661 (Table S4), much larger than its effect on physical activity study participation, 0.1826. From the EA effects alone, the ascertainment bias would appear to be much stronger with dietary study invitation. By contrast, on the genetic level, the ascertainment bias for physical study participation is stronger than that for invitation after adjustment for EA. Thus, while the genetic component to participation is associated with EA, its effects on other traits is not manifested mainly through EA. Importantly, phenotypes known to correlate with participation, do not fully capture the nature and magnitude of ascertainment bias. The estimated effects of EA are substantially larger than that of the *pPGS*. However, the *pPGS* only captures a small fraction of the full genetic component of participation, the latter could have effects that are at least comparable to those of EA, particularly for the participation traits. When males and females are analysed separately for the traits/variables in Table 1 (Fig. 2), no significant difference is found for the estimated effects of *pPGS*. Furthermore, while it has been reported that UKBB participation rates differ by sex and age^17^, the *pPGS* is associated with neither (*P* > 0.05). This implies that the effect of the *pPGS*, based on autosomal variants, is additive to the effects of sex and age on participation, with no detectable statistical interactions.

**Figure 2:**
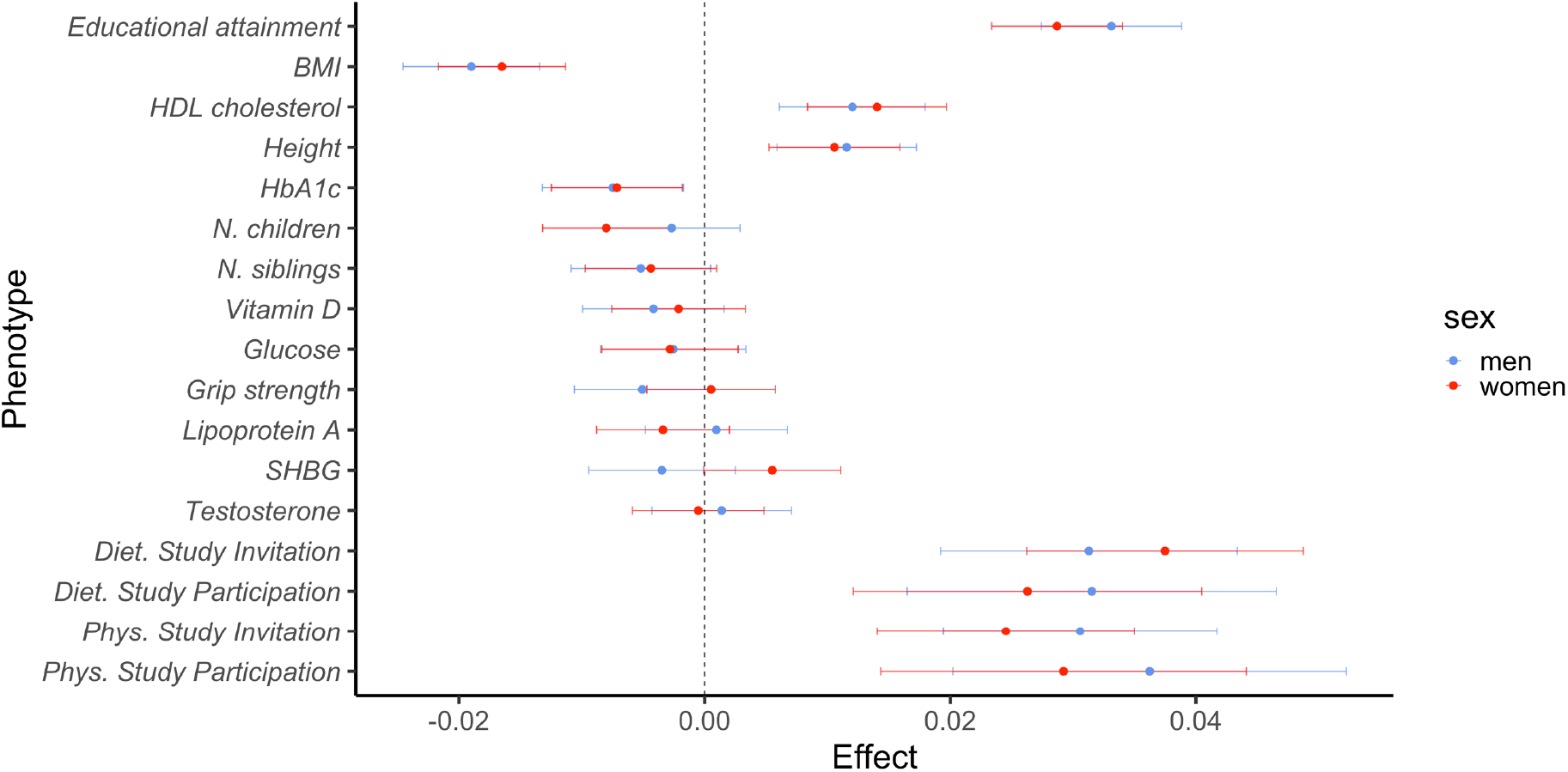
Participation PGS and sex-specific analysis. This figure depicts the results from regressing the participation polygenic score (PGS) on a range of phenotypes for men and women of White British descent who have no close relatives in UK Biobank (>3^rd^ degree for all pairs). Each dot depicts the effect in SD or log(OR) of the participation PGS (x-axis) on the corresponding phenotype (y-axis). The vertical lines depict 95% confidence intervals taking the corresponding LD-score regression intercept into account (Supp. Text). The blue dots correspond to regression in the subset of men while the red dots correspond to regression in the subset of women. Genotyping array (BiLEVE or Axiom) was an additional covariate in the regressions. For men, the sample size ranged from 111,219 individuals to 127,186 individuals and for women the sample size ranged from 124,741 to 145,223. SNP-wise weights for the PGS were computed using IBD information from siblings and parent-offspring pairs (not sexspecific weights). Prior linear regression for each sex, the quantitative phenotypes were adjusted for year of birth (YOB), age at measure and 40 PCs separately for each sex and then the residuals were standardised also separately for each sex (Supp. Text). For the binary phenotypes, we performed a logistic regression separately for each sex, including YOB, age at measure up to the order of three, 40 PCs, and genotyping array as additional covariates. Grip strength was additionally adjusted for height. BMI: body max index. HDL cholesterol: High-density lipoprotein cholesterol. HbA1c: Glycated haemoglobin. N. children: Number of children. N. siblings: Number of full siblings. SHBG: Sex hormone binding globulin.

UKBB did not recruit families and participants were all adults providing their own consent^16^. Under these conditions, for alleles that have an effect on participation, relative allele frequencies in different groups of individuals and genetic segments depend on many factors, most important of which are the overall participation rate and the participation rate of those with a close relative among the participants (Supp. Text and Fig. S9). For UKBB, ignoring confounding factors, by simulations, we estimated that the allele frequency differences between the shared and not-shared in sibling pairs are about 90% of the frequency differences between the alleles in participants than non-participants (Supp. Text). Alleles in the unrelateds are by definition not IBD shared through a recent common ancestor with any other participant. By comparison, participants with close relatives among the participants have both shared and not-shared alleles. Thus:

> The Second Principle --- On average, participants with close relatives among the other participants are enriched with alleles that promote participation relative to other participants.

To examine this, we randomly partitioned the sibling pairs into two halves, derived a *pPGS* based on a GWAS performed with the first half (*pPGS_1_*), and compare the *pPGS_1_* values of sibpairs in the second half with the unrelateds. The splitting reduces power. Nonetheless, the second half of the sibpairs have average *pPGS_1_* that is 0.041 standard deviation (SD) higher than that of the unrelateds (*P* = 2.1 × 10^-5^). Switching the roles of the two halves of the sibpairs, the first half of the sibpairs have average *pPGS_2_* that is 0.030 SD higher than that of the unrelateds (*P*= 2.0 × 10^-3^).

> The Third Principle --- If genetics contribute to participation, there would be more close relative pairs among the participants than what is expected if participation is random.

This is true because a participant has a higher than average expected participation genetic component value, and thus so would its close relatives. Even though UKBB did not recruit families on purpose, the dataset contains more close relatives (≤3^rd^ degree) than what would be expected from random sampling, *e.g*. sibling pairs are twice as many as expected^16^. They speculated that the cause of overrepresentation of close relatives is mutual consultation and possibly shared environment leading to correlated participation^16^. Here we show that shared genetics most likely also play a substantial role.

## Discussion

Ascertainment bias, particularly participation bias, is arguably the most challenging problem in applied statistics and it is becoming increasingly relevant in the age of ‘Big Data’^1,3^. In addition to obvious pitfalls with data analyses that ignored ascertainment bias, for genetic studies, more subtle consequences include collider bias^20^ that decreases correlations between contributing alleles and the introduction of artificial epistatic effects (Supp. Text). Here we show that ascertainment bias leaves many footprints in the genetic data of the participants. Exploiting that, we are able to perform a GWAS on participation using only genetic data of the participants. Notably, each individual often harbours both ‘case’ and ‘control’ alleles, at different sites and the same sites. With nearly perfectly matched controls, our GWAS only captures direct effects and is unaffected by population stratification confounding. While the genetic component to participation can only be studied through a genetic study, the former can play a role in all sampling-based studies, genetic or otherwise.

One of the complications of studying the genetics of participation is the issue of replication. As study design, participation rate and the contribution of genetics can vary hugely from study to study, each study has to be evaluated on its own and the participation GWAS results cannot be expected to replicate across all studies. For this reason, we validated our participation GWAS through a special form of within-study replication. This was done by investigating the relationship between the *pPGS* and various phenotypes in a group of UKBB participants that do not overlap with the UKBB first-degree relatives that the participation GWAS is based on. Notably, the *pPGS* is constructed from a GWAS performed without phenotypes. Thus, its associations with phenotypes such as EA and BMI, that have previously been reported to exhibit ascertainment bias^17,20^, is confirmation that our GWAS is indeed capturing the genetic effects of participation.

The results of this study demonstrate that there can be a common component that underlies many different participation events. The *pPGS* we constructed is based on information about primary participation, but it plays a role in both the passive (being invited) and active (deciding to participate when invited) phases of the secondary participation events. Thus, the relevance of a genetic component underlying participation should not be judged based on an individual selection event only. Its effect could accumulate through its impact on many participation events of a person’s lifespan, or it can be magnified through nested participation, *e.g*.the participants in the dietary and physical activity studies have higher average genetic propensity to participate than the other UKBB participants, who already have higher average propensities than the population. In contrast to a common tendency to think of participation as a consequence of other characteristics and established traits, we propose that the propensity to participate in a whole range of events should be studied as a behavioural trait in its own right. A better understanding of this behaviour could lead to better recruitment schemes, for studies genetic and otherwise, that increases participation rate and have lower ascertainment bias.

## Acknowledgements

A.K. is supported by the Li Ka Shing Foundation. S.B. is supported by the Li Ka Shing Foundation and the Goodger and Schorstein scholarship. This research has been conducted using the UK Biobank Resource under application number 11867. The computational aspects of this research were supported by the Wellcome Trust Core Award Grant Number 203141/Z/16/Z and the NIHR Oxford BRC. The views expressed are those of the authors and not necessarily those of the NHS, the NIHR or the Department of Health.

## Author contributions

Idea: A.K.

Methodology: A.K. and S.B.

Design of study: A.K. and S.B.

Investigation: A.K. and S.B.

Data application: A.K and S.B.

Writing – original draft: A.K. and S.B.

Writing – review and editing: A.K. and S.B.

## Supplementary Materials

Figs. S1 to S9

Tables S1 to S4

Supplementary Text

