## Supplementary Materials for "The Genetics of Participation: Method and Analysis"

### Supplementary Material

Stefania Benonisdottir[1,*] and Augustine Kong[1,*]

[1]Big Data Institute, Li Ka Shing Centre for Health Information Discovery, University of Oxford, Oxford, UK.

Vitamin D ... 29
Glucose ... 30
Grip strength ... 30
Lipoprotein A ... 30
Sex hormone binding globulin (SHBG) ... 31
Testosterone ... 32
Dietary study invitation ... 32
Dietary study participation ... 33
Physical activity study invitation ... 33
Physical activity study participation ... 33
**GWAS summary statistics ... 33**
**LD score regression intercepts ... 34**
**Relative allele frequencies in different groups of individuals and segments ... 34**
**Significant associations in the MHC region ... 38**
**The effects of selection on the associations between trait and genotypes ... 39**
**References ... 40**

### Supplementary Figures and Tables

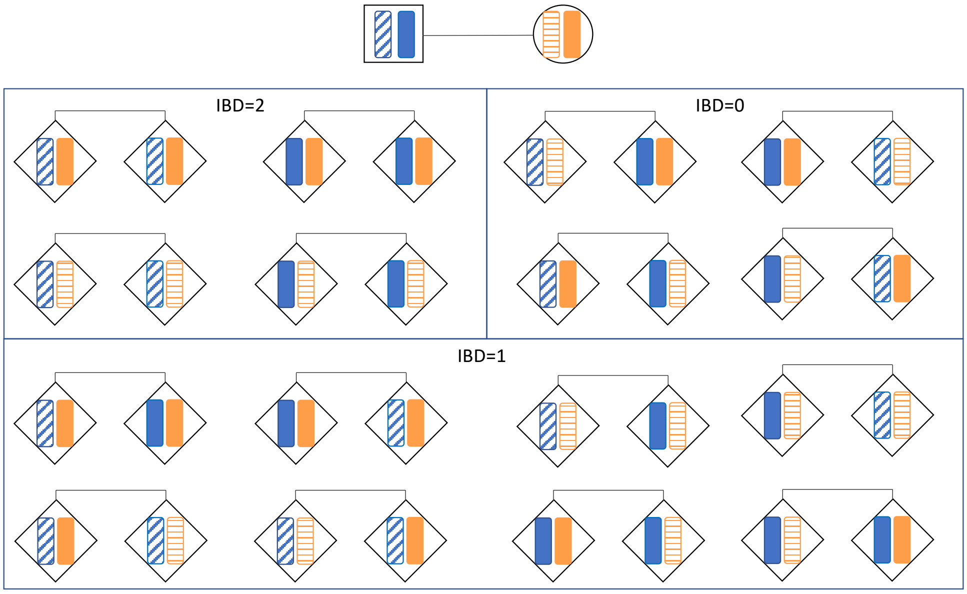

**Figure S1. Parental transmission to sibling pairs.**

Figure showing the 16 possible combinations of ordered parental genetic segments for a sibling pair at a given locus. The father is shown as a rectangular shape with two blue distinct genetic segments and the mother is shown as a circular shape with two orange distinct genetic segments. Here, distinct refers to origin. That is, the four parental genetic segments could be identical by state but they are distinct with regard to grandparental origin. The sibling pairs are shown as diamond shapes, each carrying one blue genetic segment inherited from the father and one orange genetic segment inherited from the mother. The siblings share both segments identical by descent (IBD) for ¼ of the combinations and the siblings share none of the segments IBD for ¼ of the combinations. The siblings share one segment IBD for half of the combinations. For the IBD=1 combinations, the shared segment is paternally inherited for half of them and maternally inherited for the other half.

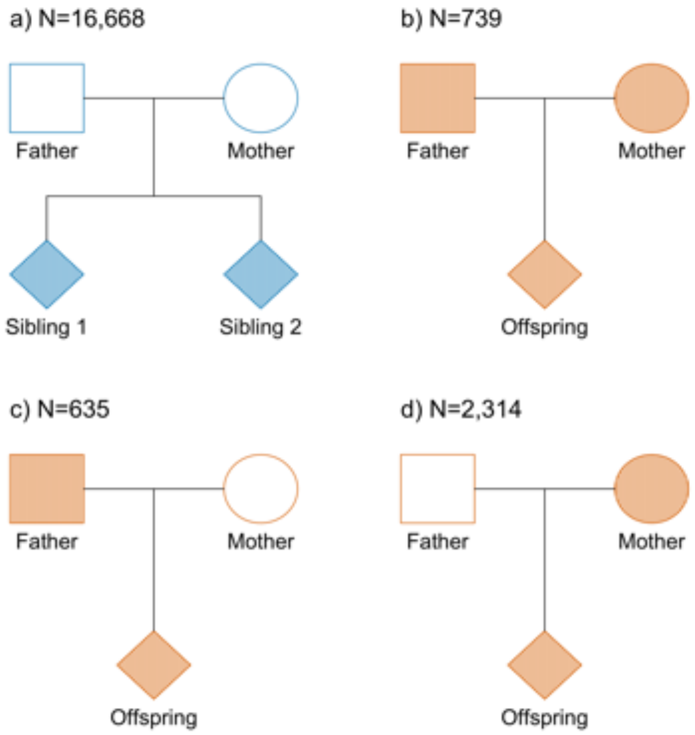

**Figure S2. First degree relatives.** Figure showing the number of first-degree relatives in UKBB included in our analysis. Shaded symbols indicate available genotype information. The 739 trios in b) count as 1,478 parent-offspring pairs and therefore the parent-offspring pairs are 4,427 in total. The groups of siblings and parent-offspring pairs do not overlap. That is, no parent or offspring in b), c) and d) have a sibling present in the dataset.

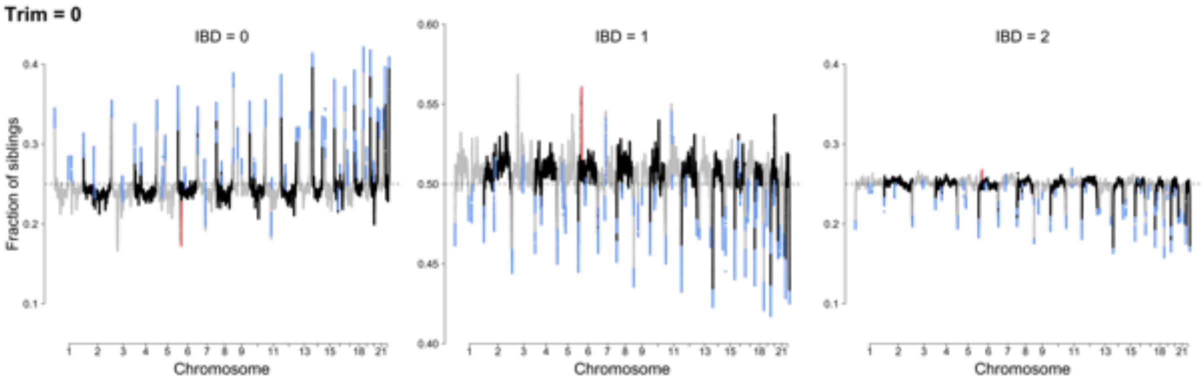

**Figure S3. Fraction of sibling sharing SNPs IBD.** For each SNP we counted how many sibling-pairs shared the SNP IBD=0, IBD=1 and IBD=2 and then divided that number with the total number of sibling pairs with non-missing values for the SNP in question. The y-axis corresponds to this fraction of siblings and the x-axis denotes the chromosomal position of each SNP. The grey dotted lines denote the expected fraction for each IBD scenario. KING inferred IBD segments for 40 distinct regions. The blue dots denote the first and last 250 SNPs in each of those 40 regions. The red dots denote variants in the MHC region (chr6:25,000,000-33,500,000). These computations are based on non-trimmed data (Trim=0). That is, SNPs in the beginning and end of IBD regions had not been trimmed away.

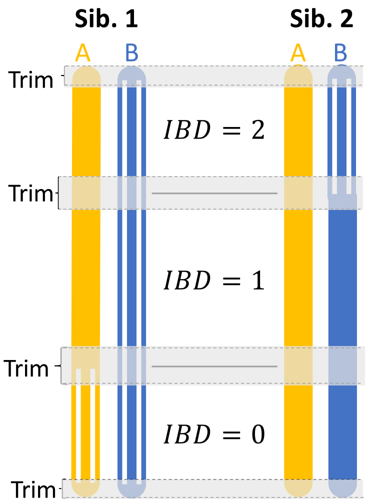

**Figure S4. Trimming SNPs at the beginning and end of IBD regions.** To account for the error-rate of inferring IBD state being higher in the beginning and end of IBD regions, we trimmed away 250 SNPs from the beginning and end of each IBD specific regions for each sibling pair.

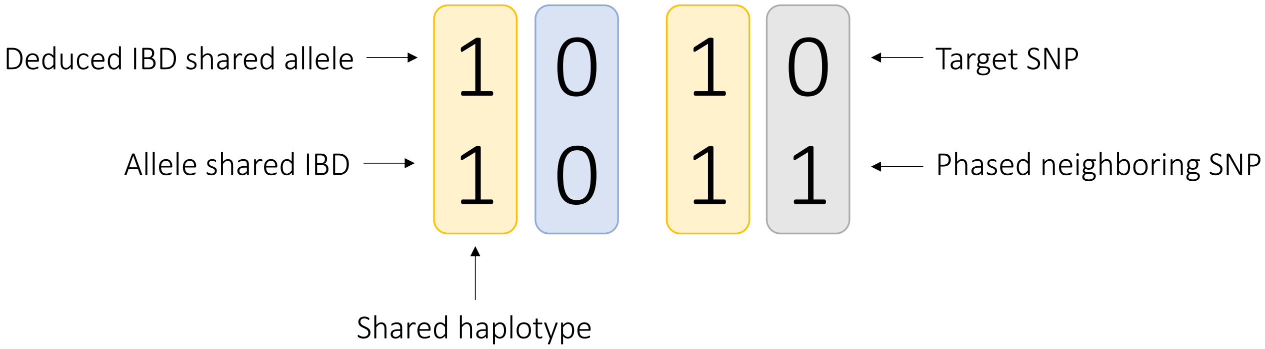

**Figure S5. Inferring shared allele for IBD=1 when both individuals are heterozygous for target SNP.** A neighbouring SNP, for which one individual is heterozygous while the other is homozygous, is informative of which allele is shared for the target SNP, for which both individuals are heterozygous, if those two SNPs are phased together.

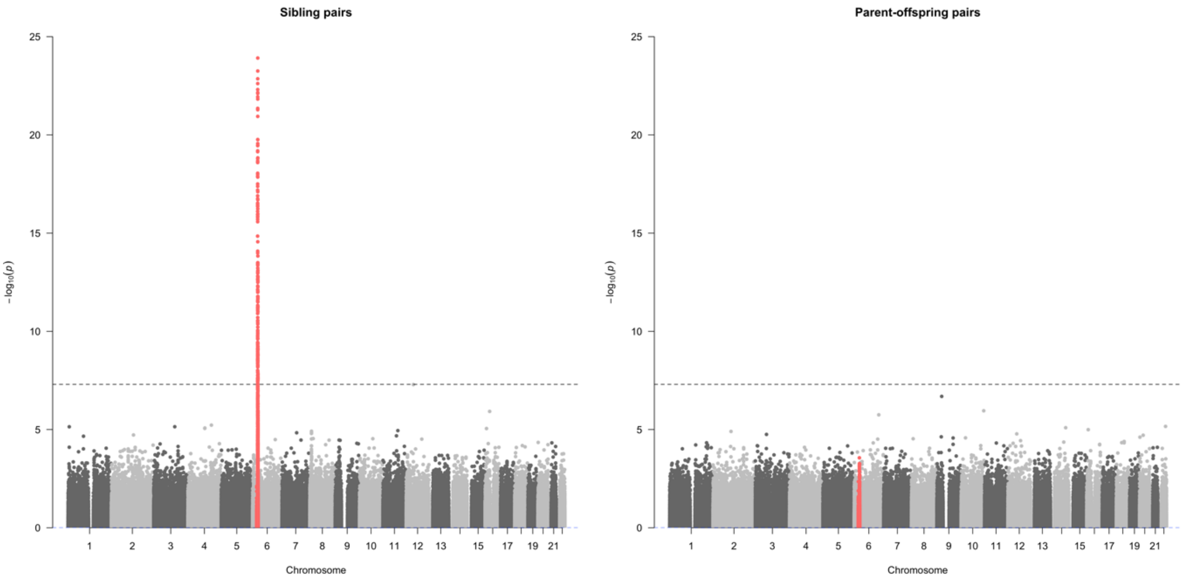

**Figure S6: Association of MHC SNPs with ascertainment: Sibling pairs vs. parent-offspring pairs.** Figure showing -$\log_{10}$(P) values (y-axis) for association with ascertainment in the group of 16,668 sibling pairs (figure on the left) and 4,427 parent-offspring pairs (figure on the right). Each dot represents a single variant association. The x-axis depicts chromosomal position in build 37. SNPs in the MHC region (chr6:25,000,000-33,500,000) are coloured red in the figure. This figure includes SNPs in extended LD regions which were otherwise excluded from our participation GWAS (see *Variant filtering* in Supplementary text). The black dotted line is set at $P$=5E-8.

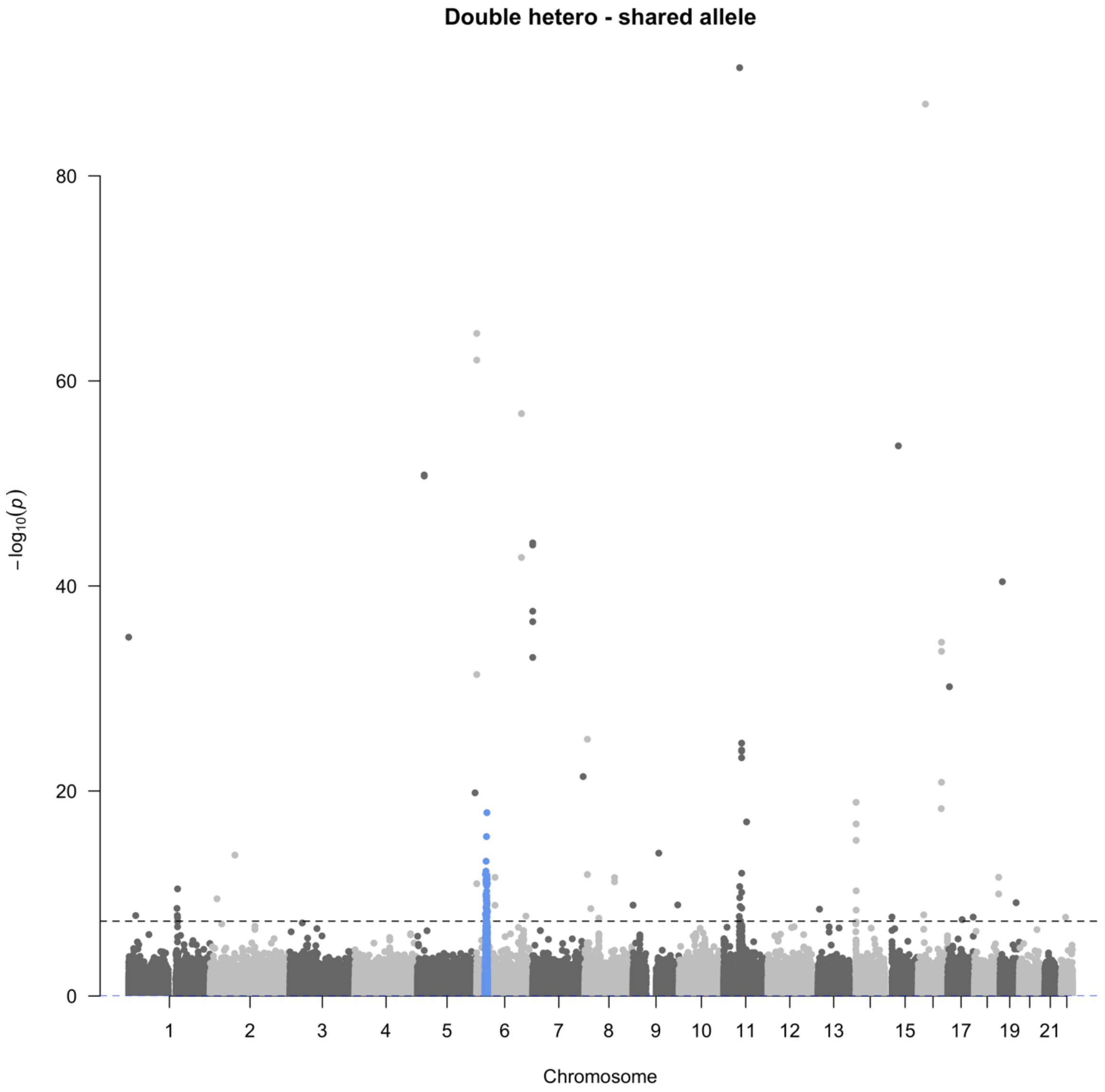

**Figure S7. Deviations from the expected fraction of the shared allele.** For each SNP, we tested if the expected fraction of the shared allele deviated from the observed fraction of the shared allele (inferred from phasing information) in the group of parent-offspring pairs that were heterozygous for the SNP in question. Each dot in the figure represents one SNP. SNPs in the MHC region, (chr6:25,000,000-33,500,000), are coloured blue in the figure. The y-axis shows $-\log_{10}(P)$ from a two-sided binomial test and the x-axis shows chromosomal position in build 37. The black dotted line is set at *P*=1.0E-6.

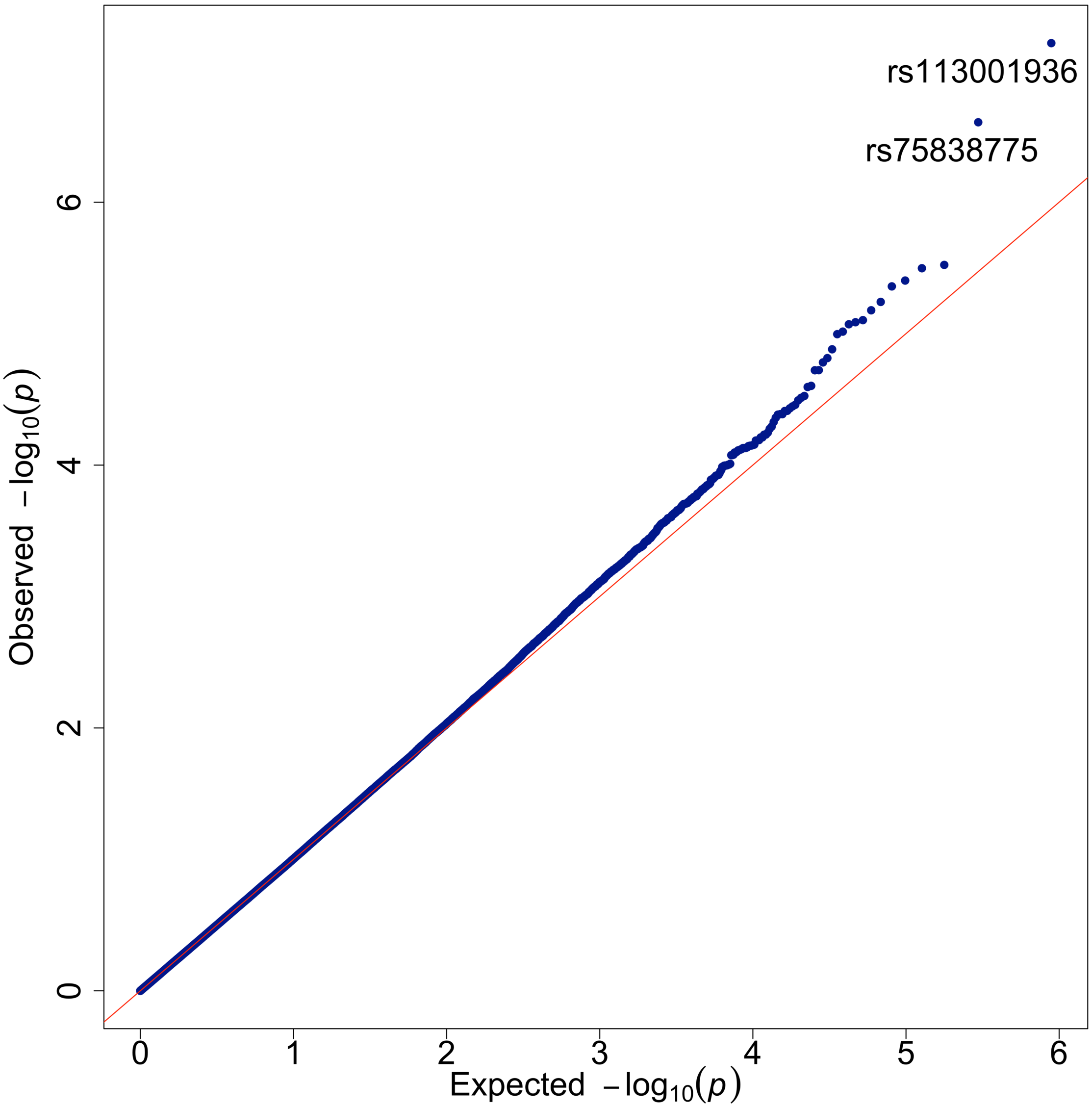

**Figure S8. QQ-plot for the Participation GWAS.** This figure shows the expected -$\log_{10}$P-values and the observed -$\log_{10}$P-values from the GWAS comparing shared and not-shared alleles of first degree relatives in UK Biobank. We restricted our analysis to 444,457 high-quality SNPs in the phasing input that had MAF>1% and fulfilled various quality criteria (see *Variant filtering* in Supplementary Text).

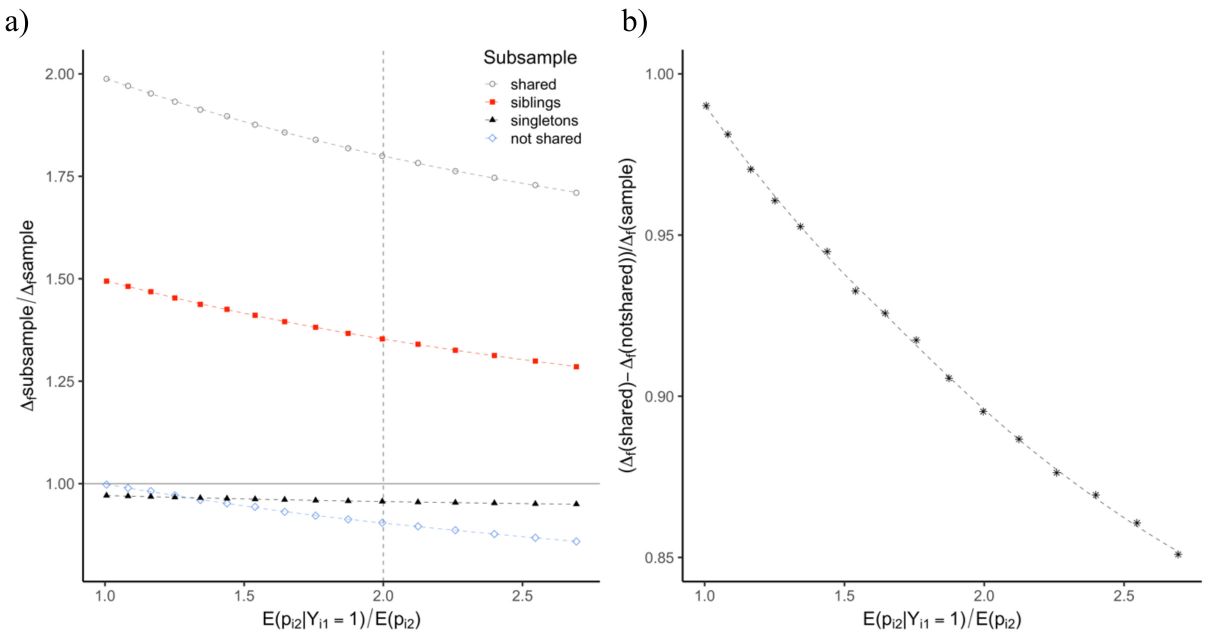

**Figure S9. Relative frequency differences as a function of enrichment of sibling pairs in sample.** In both figures the x-axis denotes induced enrichment of sibling pairs in sample. $E(p_{i2})$ stands here for the expected participation probability for sibling 2 of sib-pair $i$ while $E(p_{i2}|Y_{i1} = 1)$ is the expected participation probability of sibling 2 given that sibling 1 participates. The figures show results from simulations based on a model where the participation probability for sibling $j = \{1, 2\}$ of pair $i$ is described with the logistic regression model, $logit(p_{ij}) = \alpha + \beta X_{ij}$, with $X_{ij} = w_1 G_{ij} + w_A A_i + w_B B_{ij}$. Here, $G_{ij}$ denotes a standardized genotype for a single *SNP* and $A_i$ and $B_{ij}$ are standard normally distributed variables. All three variables, $G_{ij}$, $A_i$ and $B_{ij}$, were assumed to be independent of each other. We assumed that $w_1^2 + w_A^2 + w_B^2 = 1$ so that $var(X_{ij}) = 1$. $B_{i1}$and $B_{i2}$ were assumed to be independent, hence the correlation between $X_{i1}$ and $X_{i2}$ was induced by the shared component $A_i$ and the correlation between $G_{i1}$ and $G_{i2}$. For the simulations, we chose $\alpha = -3.48$ and $\beta = 1.25$. This ensured that $E(p_{ij}) \approx 0.055$. For $\pi =$ $0.0, 0.1, 0.2, 0.3, 0.4, 0.5, 0.6, 0.7, 0.8, 0.9, 1.0, 1.1, 1.2, 1.3, 1.4,$ and $1.496$, we simulated 500 replications from a population of $10^8$ sib-pairs with $w_1^2 = \pi \cdot 0.01$ , $w_A^2 = \pi\left(\frac{2}{3} - 0.005\right)$ and $w_B^2 = 1 - \pi\left(\frac{2}{3} + 0.005\right)$. **Figure a)** depicts how the difference between subsample allele frequency and population allele frequency, $\Delta_f subsample$, divided by the difference between total sample allele frequency and population allele frequency, $\Delta_f sample$, (y-axis) is a function of the induced enrichment of sibling pairs in sample (x-axis). Each dot represents a mean for the 500 replications for a given $\pi$ and subsample. The subsamples are as following: 1) shared alleles of participating sibling pairs (grey circles), 2) alleles of participating siblings (red squares), 3) alleles of singletons, that is participants with a non-participating sibling (black triangles) and 4) not-shared alleles of participating siblings (blue diamonds). **Figure b)** depicts how the difference between the frequency of shared alleles and not shared alleles of participating siblings, $\Delta_f shared - \Delta_f notshared$, divided by the difference between total sample allele frequency and population allele frequency, $\Delta_f sample$, (y-axis) is a function of the induced enrichment of sibling pairs in sample (x-axis). Each dot in the figure represents a mean for the 500 replications for a given $\pi$.

| | | *Second participating sibling* | |
|---|---|---|---|
| | | **Brother** | **Sister** |
| *First participating sibling* | **Brother** | 3,307 | 3,920 |
| | **Sister** | 3,593 | 5,848 |

**Table S1. Brother-sister division.** Table showing how the 16,668 sibling pairs are divided into brother-brother, sister-sister, brother-sister and sister-brother pairs. Given that the male fraction is 43.4% among first participating siblings and 41.4% among second participating siblings, the fraction of brother-sister pairs is lower than what we would expect if we were to randomly draw pairs from two infinite samples with the same male-female fractions ($P=5.8\times10^{-17}$, chi-square test).

| Chromosome | BP start | BP end |
|---|---|---|
| 1 | 48,000,000 | 52,000,000 |
| 2 | 86,000,000 | 100,500,000 |
| 2 | 134,500,000 | 138,000,000 |
| 2 | 183,000,000 | 190,000,000 |
| 3 | 47,500,000 | 50,000,000 |
| 3 | 83,500,000 | 87,000,000 |
| 3 | 89,000,000 | 97,500,000 |
| 5 | 44,000,000 | 51,500,000 |
| 5 | 98,000,000 | 100,500,000 |
| 5 | 129,000,000 | 132,000,000 |
| 5 | 135,500,000 | 138,500,000 |
| 6 | 25,000,000 | 33,500,000 |
| 6 | 57,000,000 | 64,000,000 |
| 6 | 140,000,000 | 142,500,000 |
| 7 | 55,000,000 | 66,000,000 |
| 8 | 8,000,000 | 12,000,000 |
| 8 | 43,000,000 | 50,000,000 |
| 10 | 37,000,000 | 43,000,000 |
| 11 | 45,000,000 | 57,000,000 |
| 11 | 87,500,000 | 90,500,000 |
| 12 | 33,000,000 | 40,000,000 |
| 12 | 109,500,000 | 112,000,000 |
| 20 | 32,000,000 | 34,500,000 |

**Table S2. Extended LD regions.** Regions of extended linkage-disequilibrium (LD) as reported by Bycroft et al. 2018 [1]. Base pair (BP) positions refer to build 37.

| | Dietary Study | Physical Activity Study | Both studies |
|---|---|---|---|
| Total | 272,409 | 272,409 | 272,409 |
| Invited | 166,993 | 132,633 | 107,061 |
| Participated | 54,124 | 59,455 | 23,027 |

**Table S3: Secondary participation.** This table depicts the numbers of invitees and participants in two follow up studies in UK Biobank 1) The Dietary Study and 2) The Physical Activity Study. The numbers shown here apply to White British individuals that do not have any close relatives within UK Biobank (>3rd degree for all pairs)

| Phenotype | | | Adjusted for pPGS | | |
|---|---|---|---|---|---|
| **Quantitative traits** | **Effect** | **P-value** | **Effect** | **P-value** | **N** |
| AFB | 0.3500 | $< 1.0 \times 10^{-300}$ | 0.3494 | $< 1.0 \times 10^{-300}$ | 95,424 |
| BMI | -0.1395 | $< 1.0 \times 10^{-300}$ | -0.1390 | $< 1.0 \times 10^{-300}$ | 260,118 |
| HDL cholesterol | 0.0993 | $< 1.0 \times 10^{-300}$ | 0.0990 | $< 1.0 \times 10^{-300}$ | 227,964 |
| Height | 0.1458 | $< 1.0 \times 10^{-300}$ | 0.1456 | $< 1.0 \times 10^{-300}$ | 260,389 |
| HbA1c | -0.0728 | $1.1 \times 10^{-272}$ | -0.0726 | $3.3 \times 10^{-271}$ | 248,889 |
| Nchildren | -0.0550 | $3.2 \times 10^{-170}$ | -0.0549 | $2.4 \times 10^{-169}$ | 260,070 |
| Nsib | -0.1454 | $< 1.0 \times 10^{-300}$ | -0.1454 | $< 1 \times 10^{-300}$ | 256,956 |
| Vitamin D | -0.0283 | $9.8 \times 10^{-43}$ | -0.0283 | $1.5 \times 10^{-42}$ | 238,755 |
| Glucose | -0.0260 | $1.5 \times 10^{-34}$ | -0.0259 | $2.1 \times 10^{-34}$ | 227,812 |
| Grip strength | 0.0468 | $2.1 \times 10^{-121}$ | 0.0469 | $7.9 \times 10^{-122}$ | 259,141 |
| Lipoprotein A | -0.0099 | $1.7 \times 10^{-6}$ | -0.0099 | $1.9 \times 10^{-6}$ | 241,318 |
| SHBG | 0.0348 | $9.1 \times 10^{-59}$ | 0.0348 | $1.4 \times 10^{-58}$ | 226,220 |
| Testosterone | 0.0078 | $1.1 \times 10^{-4}$ | 0.0078 | $1.1 \times 10^{-4}$ | 246,979 |
| **Binary traits** | **log(OR)** | **P-value** | **log(OR)** | **P-value** | **Ncases/Ncontrols** |
| Dietary study invitation | 0.4661 | $< 1.0 \times 10^{-300}$ | 0.4655 | $< 1.0 \times 10^{-300}$ | 159,956/100,994 |
| Dietary study participation | 0.2621 | $< 1.0 \times 10^{-300}$ | 0.2615 | $< 1.0 \times 10^{-300}$ | 51,891/108,065 |
| Physical activity study invitation | 0.3262 | $< 1.0 \times 10^{-300}$ | 0.3256 | $< 1.0 \times 10^{-300}$ | 127,024/133,926 |
| Physical activity study participation | 0.1826 | $2.5 \times 10^{-212}$ | 0.1818 | $1.7 \times 10^{-210}$ | 56,817/70,207 |

**Table S4: Educational attainment and association with phenotypes.** This table depicts how educational attainment in terms of years of education is associated with a range of phenotypes. For the quantitative traits, we performed linear regression with the corresponding phenotype as a response and educational attainment and genotyping array (BiLEVE or Axiom) as explanatory variables in the subset of individuals of White British descent who have no close relatives in UK Biobank (>3rd degree for all pairs). Prior to regression analysis, the quantitative traits, including educational attainment, were adjusted for year of birth (YOB), age at measure and 40 PCs separately for each sex and then the residuals were standardised also separately for each sex (see Supp. Text). Grip strength was additionally adjusted for height. BMI: body max index. HDL cholesterol: High-density lipoprotein cholesterol. HbA1c: Glycated Haemoglobin. Nchildren: Number of children. Nsib: Number of full siblings. SHBG: Sex hormone binding globulin. Information about age at first birth (AFB) was only available for women. For the binary phenotypes, we performed a logistic regression including YOB, age at measure up to the order of three, 40 PCs, sex and genotyping array as additional covariates. Columns 2 and 3, labelled 'Effect/log(OR)' and 'P-value', depict the slope/log(odds-ratio) for educational attainment and the corresponding P-value. The P-values have been adjusted with the LD-score regression intercept of the corresponding phenotype. Columns 4 and 5 show the slope/log(OR) and the P-value for educational attainment when study participation PGS (pPGS) has been added as an additional covariate. SNP-wise weights for the study participation PGS were computed using identity-by-descent information from 16,668 siblings and 4,427 parent-offspring pairs in UK Biobank.

### Supplementary Text

#### Genotype Data

*Individuals*

The UK Biobank (UKBB) is a prospective cohort that includes approximately 500 thousand genotyped and phenotyped individuals from across the United Kingdom (UK)[1]. We restricted our analysis to 402,199 individuals in the UKBB who had not withdrawn consent, did not have a duplicate/twin in sample and fulfilled the following critera in the sample quality control file provided in the UKBB data, 'ukb_sqc_v2.txt': (1) Did not have excess of third degree relatives, (2) was not a heterozygosity or missingness outlier (3), had missing rate below 2%, (4) was included in the kinship inference, (5) was included in the phasing input, (6) did not show potential sex chromosome aneuploidy, (7) did not demonstrate potential sex mismatch and (8) had been determined to be White British (WB) according to self-identified ancestry and PCA[1].

*SNPs*

Our analysis was restricted to 658,722 directly-genotyped SNPs that were available in the UKBB phased haplotype data[1]. The genotyping process and quality control for the SNPs in the phasing input as well as the phasing process was described in detail by Bycroft et al (2018)[1]. Briefly, SHAPEIT3 was used to attain phased haplotypes for directly-genotyped SNPs that fullfilled the following criteria: 1) Present on both the UK BiLEVE and UK Biobank Axiom array, 2) missing rate <5%, 3) MAF>0.0001, 4) did not demonstrate batch effect, 5) did not demonstrate array effect, 6) did not demonstrate discordance across control replicates and 7) did not fail quality control, (plate effect, Hardy-Weinberg equilibrium (HWE) and sex effect), in more than one of 109 batches of ~4,900 individuals.

**Identifying relatives**

The UKBB data includes kinship coefficients, attained with the program KING[2], for pairs that were genetically inferred to be related of 3rd degree or closer. Among those pairs, we identified first-degree relatives of WB descent, 16,668 sibling pairs and 4,427 parent-offspring pairs, using kinship coefficient boundaries recommended by the authors of KING[2]. Pairs with kinship coefficient >0.177 and <0.354 were concluded to be first degree relatives and, following Bycroft et al. (2018)[1], those who shared less than 0.12% IBD=0, were classified as parent-offspring pairs

These two groups, siblings and parent-offspring pairs, did not overlap. That is, none of the sibling pairs had a parent present in the dataset and none of the parent-offspring pairs had a sibling present in the dataset (Fig. S2). The parent-offspring pairs were made up of 739 trios, 635 father-offspring pairs and 2,314 mother-offspring pairs. The age-difference between the parents and their offspring ranged from 15 to 31 years with the mean being 23.43 years (SD=2.71).

For sibling-groups with more than 2 siblings in the dataset, we chose the first two participating siblings in the group. The mean year of birth of first participating siblings was 2 months lower than the mean year of birth of the second participating siblings. Given that the male fraction was 43.4% among first participating siblings and 41.4% among second participating siblings, the fraction of brother-sister pairs (45.1%) was lower than what we would expect if we were to randomly draw pairs from two infinite samples with the same male-female fractions ($P=5.8\times10^{-17}$, chi-square test, Table S1).

Within the WB subset, 272,409 individuals had no close relatives within the UKBB (>3$^{rd}$ degree for all pairs). We refer to this set of individuals as the 'unrelated' set. The male fraction was 46.7% in the 'unrelated' set compared to 31.0% among the parents and 42.4%

among the siblings. In addition, the ‘unrelated’ individuals were on average 5.6 months younger than the sibling pairs.

**Inferring IBD segments of sibling pairs**

Parent-offspring pairs share all SNPs IBD=1, unless the offspring harbours de-novo mutations. Sibling pairs can share 0, 1 or 2 alleles IBD with the expected fraction of SNPs being 0.25 for IBD=0, 0.5 for IBD=1 and 0.25 IBD=2. We inferred IBD segments for sibling pairs using the option --ibdseg in the program KING (version 2.2.4)[2]. The authors of KING have not yet published a manuscript describing their IBD segment algorithm but claim in their manual that it has been well tested (http://people.virginia.edu/~wc9c/KING/manual.html, accessed September 8th 2020). The IBD-segments were inferred from genotypes of 649,509 SNPs that conformed to HWE in a subgroup of 272,234 WB individuals in the ‘unrelated’ set ($P>1.5\times10^{-23}$, exact test). HWE was tested with the --hardy option in PLINK 1.90[3]. Before inferring IBD, KING divided the chromosomes into 40 regions but 54 SNPs were not included in those regions and were therefore excluded from subsequent analysis.

**Inferring shared allele**

For a given biallelic SNP and a pair of first-degree relatives, there are 5 possible unordered combinations of genotypes when IBD=1. As IBD implies identical by status (IBS), it is trivial to infer what allele is shared IBD when both individuals are homozygous for the same allele, or when one is heterozygous and the other is homozygous. For the double heterozygous scenario, we used the phasing information provided by UKBB to infer which allele was shared. For a target SNP that both first degree relatives were heterozygous for, we used the closest neighbouring SNP, for which one individual was heterozygous while the other was homozygous, to determine which strand was shared and subsequently which allele of the target SNP was shared (Fig. S5).

**Participation GWAS**

*Test statistics*

By comparing shared and not-shared alleles of first-degree relatives in UK Biobank we computed the following three t-statistics for directly genotyped SNPs available in the phased genotype data:

*DPO: T-statistic* comparing the allele frequency of the SNP in question for shared and not-shared alleles of the 4,427 WB parent-offspring pairs.

*DSIB1: T-statistic* comparing the allele-frequency of shared and not-shared alleles of the sibling pairs sharing the SNP in question IBD=1.

*DSIB20: T-statistic* comparing the allele frequency in the group of siblings sharing the SNP in question IBD=2 to the allele frequency in the group of siblings sharing the same SNP IBD=0.

The sample size for the sibling IBD specific allele frequencies varied between variants as we trimmed 250 SNPs in the beginning and end of each IBD region for each sibling pair (see *IBD trimming* below). In the subset of 444,457 variants that passed various quality control thresholds (see *Variant-filtering* below) the sample size for sibling IBD=1 allele frequencies ranged from 6,861 siblings to 8,242 siblings (mean=7,683; SD=220), the sample size for IBD=0 allele frequencies ranged from 2,841 siblings to 4,102 siblings (mean=3,613; SD=120) and the sample size for IBD=2 allele frequencies ranged from 3,169 siblings to 4,102 siblings (mean=3,766; SD=158).

*Variant-filtering*

We restricted our GWAS results to 444,457 high-quality SNPs that fulfilled the following criteria:

- Conformed to HWE (P>0.001, exact test, PLINK 1.90b[3]) in a group of 272,234 WB individuals with no close relatives in UK Biobank.
- Not among the 250 SNPs in the beginning and end of each of the 40 regions that KING inferred IBD for.
- Not in extended linkage disequilibrium (LD) regions (Table S2).
- With MAF>1% in the set of 16,668 sibling pairs
- With a rsname
- With non-ambiguous nucleotides
- Conformed to shared allele expectations in the group of double heterozygous parent-offspring pairs ($P>1.0\times10^{-6}$, two-sided binomial test, see *Phasing criteria* below).
- Conformed to IBD fraction expectations for sibling pairs (see *IBD fraction criteria* below).

*Phasing criteria*

For each SNP, we investigated the accuracy of the phasing provided by UKBB[1] by comparing the expected fraction of the shared allele to the observed fraction of the shared allele in the group of parent-offspring pairs that were heterozygous for the SNP in question. A heterozygous parent passes either allele, 0 or 1, with an equal probability to his/her offspring. If the offspring is heterozygous as well, the probability of the shared allele being the one coded as 1 is simply the probability of the allele inherited from the other parent being the one coded as 0. Assuming random mating, the allele inherited from the other parent can be regarded as a randomly drawn allele from the population and hence, in the double heterozygous scenario, the probability of the shared allele being the one coded as 1 is the population allele frequency of the allele coded as 0. We removed 262 SNPs for which the observed fraction of the shared allele, among the double heterozygous parent-offspring pairs , deviated from the expected fraction at a threshold of $P<1.0\times10^{-6}$ (two-sided binomial test, Fig. S7). Of those 262 SNPs, 85 SNPs deviated from HWE with $P<0.001$ (exact test) of which 60 SNPs deviated from HWE with $P<1.0\times10^{-23}$ (exact test).

*IBD trimming*

To investigate the accuracy of the IBD segment algorithm of KING, we computed, for each SNP, the fraction of sibling pairs sharing the SNP in question IBD=0, IBD=1 and IBD=2 (Fig. S3). The deviations from expected IBD fractions were at their highest around the beginning and end of the 40 regions that KING infers IBD for (Fig. S3) consistent with the notion that the error-rate of IBD-segment placement is expected to be higher for SNPs close to the edges of IBD regions for a given sibling pair. To account for this uncertainty, we trimmed away 250 SNPs from the beginning and end of the inferred IBD-regions for each sibling pair before computing the test statistics. This trimming removed 20,000 SNPs all together (2×250 SNPs in each of the 40 regions KING infers IBD for) and reduced the sample size (number of sibling pairs) for the remaining SNPs (median reduction of 1,516 pairs).

*IBD fraction criteria*

To further account for uncertainty in the IBD inferring algorithm of KING (see above), we removed SNPs for which there was strong evidence of IBD inferring problems. In particular, we removed all SNPs for which the observed fraction of sibling pairs sharing that SNP in question IBD = 0, 1 or 2 was more than 6 standard deviations above the expected fraction,

0.25, 0.5 and 0.25 respectively. Note that if one IBD status (0, 1 or 2) is overcalled by KING then at least one of the two other IBD statuses is undercalled for that specific SNP. Hence, thresholding the IBD fractions from above also affects the lower bounds.

*Phasing error rate*

For each SNP, we estimated the phasing error rate from trios where the offspring and one parent were heterozygous while the other parent was homozygous. In particular, we defined the phasing error rate as the fraction of wrongly inferred shared allele of the double heterozygous parent-offspring pairs (from phasing information, Fig. S5) given the genotype of the homozygous parent. For the 444,457 common high-quality SNPs (see *Variant filtering* above), the average error rate was 0.0049 with the number of trios underlying the statistic for each SNP ranging from 4 to 235.

*Phasing error adjustments*

Even though the error rate in determining the shared allele through phasing when IBD = 1 and double- heterozygotes is quite low, this error can lead to a systematic bias instead of simply adding noise. This is because when the relative pairs are double heterozygous at an IBD=1 site, the minor allele is more likely to be the shared allele. In particular, denoting the frequency of allele 1 as $f$, assuming HWE and no selection, the probability that the shared allele is allele 1 conditioning on IBD=1 and double-heterozygotes is $(1 - f)$. Thus random phasing errors would lead to a positive bias in calling the major allele the shared allele. We account for this frequency ($f$) dependent bias by applying the following adjustments to the computed IBD1 shared and not shared frequencies:

$$F^*_{IBD1s} = F_{IBD1s} + \gamma \cdot n_{DH}/n_{IBD1}$$

$$F^*_{IBD1Ns} = F_{IBD1Ns} - \gamma \cdot n_{DH}/n_{IBD1}$$

$F_{IBD1s}$ and $F_{IBD1Ns}$ denote the shared and not-shared allele frequencies for pairs sharing the SNP in question IBD=1. $n_{DH}$ denotes the number of double heterozygous pairs sharing the SNP in question IBD=1 and $n_{IBD1}$ is the number of pairs sharing the SNP IBD=1. Here, $\gamma$ is a function $f$ and is determined as follows. Using the parent-offspring trios described above, for each of the 444,457 common high-quality SNPs, for cases when a parent-offspring pair is double heterozygous and the other parent is homozygous, we calculate the fraction of times the shared allele is 1 (based on the genotype of the other parent) but called 0 through phasing, $error(1|0)$, and the fraction of times when the shared allele is 0 but called 1 through phasing, $error(0|1)$. We then regress $[error(0|1) - error(1|0)]$, capturing the negative bias of calling allele 1 the shared allele, on the odd powers of the centred allele frequency, $cf = (f - 0.5)$, up to power 5. The fitted regression leads to the adjustment parameter

$$\gamma = -0.0040 \cdot (f - 0.5) - 0.0557 \cdot (f - 0.5)^3 + 0.1978 \cdot (f - 0.5)^5.$$

Note that regressing on $cf$ instead of $f$, and only the odd powers, ensures our fit/adjustment is invariant to which allele, *e.g.* major or minor, of a SNP is denoted 1. In particular, $\gamma$ is 0 for $f = 0.5$. This does not mean there is no error in calling the shared allele based on phasing. Rather, this means that when $f = 0.5$, the expected number of errors of calling 1 the shared allele when it is not is the same as the expected number of errors of calling 0 the shared allele when it is not. Also, $\gamma$ is positive for $f < 0.5$ and negative for $f > 0.5$. This is because the phasing induced bias for calling the common allele the shared allele is positive, and thus the adjustment is negative.

The phasing error adjustment was applied to the sibling shared and not shared IBD1 frequencies and the parent-offspring transmitted and not transmitted frequencies separately before computing the test-statistics.

*Major allele adjustment*

After the adjustments described above, the tendency for the major allele to be positively associated with participation was substantially reduced but not completely eliminated. To account for this, we regressed the adjusted T-statistics on centred allele frequency $(f-0.5)$ and treated the remaining residuals as the adjusted-T-statistic. The *IBD1* T-statistics were regressed on $(f-0.5)$, $(f-0.5)^3$ and $(f-0.5)^5$ while the *IBD20* T-statistics were regressed on $(f-0.5)$. Again, basing the regression on centred allele frequency, $f-0.5$, and setting the smoothing terms at the order of 3 and 5, ensured that the adjustment for a given SNP would be invariant to which allele, major or minor, was denoted as 1.

**Participation polygenic score**

Participation polygenic scores (PGS) were computed by summing over weighted alleles, using PLINK 1.90b[3], of 444,457 high-quality biallelic SNPs with MAF>1%, non-ambiguous nucleotides and fulfilled various additional quality control (see *Variant filtering* above). Each SNP was weighted by its corresponding T-statistic from the participation GWAS (see above) and the allele corresponding to a positive T-statistic was regarded as the effect-allele.

**Phenotypes**

By using various data-fields in the UKBB data release, we constructed a range of phenotypic variables for the 272,409 WB 'unrelated' individuals. The phenotypes of interest were adjusted for year of birth (data-field 34), age at recruitment (data-field 21022), genotype measurement batch (data-field 22000), and sex (data-field 31) when applicable. We accounted for population stratification by using principal components (PCs) which were inferred with the program ProPCA[4]. Details about those PCs are described in a previous publication[5]. Quantitative phenotypes were rank-based inverse normalised[6] except for educational attainment as it was not approximately normally distributed. Details about phenotype specific protocols and adjustments are provided below in the same order as the results in Table 1.

*Educational attainment*

We constructed an educational attainment variable from the data-fields ‘Qualifications’ (data-field 6138) and ‘Age completed full-time education’ (data-field 845). Both of these data-fields contain answers to questions that participants were asked on a touch-screen in the initial assessment visit. The data-field ‘Qualifications’ denotes answers to the multiple-choice question: ‘Which of the following qualifications do you have? (You can select more than one).’ We mapped the multiple-choice options to years of schooling using the ISCED classification[7] (see below), except for the option ‘NVQ or HND or HNC or equivalent’. In particular, years of schooling of NVQ/HND/HNC holders was defined to be their answer in the data-field ‘Age completed full-time education’ minus five. Individuals that answered ‘Prefer not to answer’ were mapped to ‘Not applicable’ and were excluded from subsequent analysis.

1) College or University degree: 20 years
2) A levels/AS levels or equivalent: 13 years
3) O levels/GCSEs or equivalent: 10 years
4) CSEs or equivalent: 10 years
5) NVQ or HND or HNC or equivalent: (Age completed full time education – 5) years
6) Other professional qualifications eg: nursing, teaching: 15 years
7) None of the above: 7 years
8) Prefer not to answer: Not applicable

Respondents who selected multiple options were assigned their highest category (maximum years of schooling). Individuals who had maximum year of schooling below 7 years were excluded from subsequent analysis.

In the subset of 260,950 WB 'unrelated' individuals with applicable entries for year-of-schooling, we adjusted for year of birth up to the order of three, age at recruitment and 40 PCs separately for each sex. This was done with a linear regression in R where years-of-schooling equivalent was treated as a quantitative dependent variable. The mean centred and standardised residuals from these two regressions (men and women) were used as an approximation for educational attainment in subsequent analysis.

*Age at first birth (AFB)*

The AFB variable was constructed from answers to the touchscreen question 'How old were you when you had your FIRST child?' which was asked in the initial assessment visit, (data-field 2754: 'Age at first live birth'). This information was collected from women who had indicated that they had given birth to a child in an answer to a previous question in the touchscreen questionnaire (data-field 2734: 'Number of live births'). We excluded entries below 12. In the subset of 98,653 WB 'unrelated' women with applicable AFB entries, we adjusted for year of birth up to the order of three, age at recruitment and 40 PCs by performing linear regression in R. The resulting residuals were then rank-based inverse normalised.

*Body mass index (BMI)*

The BMI variable was based on the data-field 21001-0-0: 'Body Mass index (BMI)', which was computed from height and weight measurements recorded in the initial assessment visit. In the subset of 271,535 WB 'unrelated' individuals with BMI values, we adjusted for year of birth, age at recruitment up to the order of three and 40 PCs separately for each sex by performing linear regression in R treating rank-based inverse normalised BMI values as a dependent variable. The resulting residuals, attained for males and females separately, were then rank-based inverse normalised as one group.

*High-Density-Lipoprotein (HDL) cholesterol*

The HDL cholesterol variable was based on data-field 30760-0-0: ‘HDL Cholesterol’, a biochemistry marker measured in blood samples collected at recruitment. In the group of WB ‘unrelated’ individuals, 34,628 individuals had missing HDL cholesterol values and, as indicated by data-field 30765: ‘HDL cholesterol missing reason’ and data-field 30766: ‘HDL cholesterol missing reportability’, 3 of those were missing because the values were below the reportable range and one was above the reportable range. We imputed the ‘below reportable range’ entries to the minimum reportable HDL cholesterol measurement in the group of WB ‘unrelated’ individuals minus 0.001, that is to 0.218, and the ‘above reportable range’ entry to the maximum reportable HDL cholesterol measurement plus 0.001, that is to 4.402. Then, in the group of 237,785 WB ‘unrelated’ individuals with applicable values, we adjusted the HDL cholesterol values for year of birth, age at recruitment up to the order of three and 40 PCs separately for each sex by performing linear regression in R treating rank-based inverse normalised HDL cholesterol values (including the imputed ones) as a dependent variable. The resulting residuals, attained for males and females separately, were then rank-based inverse normalised as one group.

*Height*

The height variable was based on data-field 50-0-0: ‘Standing height’, which was measured in the initial assessment visit. In the subset of 271,820 WB ‘unrelated’ individuals with height measurements, we adjusted for year of birth, age at recruitment up to the order of three and 40 PCs separately for each sex by performing linear regression in R treating rank-based inverse normalised height values as a dependent variable. The resulting residuals, attained for males and females separately, were then rank-based inverse normalised as one group.

*Glycated haemoglobin (HbA1c)*

The HbA1c variable was based on data-field 30750-0-0: ‘Glycated haemoglobin (HbA1c)’, a biochemistry marker measured in blood samples collected at recruitment. In the group of WB ‘unrelated’ individuals, 12,926 individuals had missing HbA1c values. As indicated by data-field 30755: ‘Glycated haemoglobin (HbA1c) missing reason’ and data-field 30756 ‘Glycated haemoglobin (HbA1c) reportability’, 111 of those were missing because the values were below reportable range. We imputed the ‘below reportable range’ entries to the minimum HbA1c measurement minus 0.001, that is to 15.299. In the group of 259,594 WB ‘unrelated’ individuals with applicable values, we adjusted HbA1c levels for year of birth, age at recruitment up to the order of three and 40 PCs separately for each sex by performing linear regression in R treating rank-based inverse normalised HbA1c values (including the imputed ones) as a dependent variable. The resulting residuals, attained for males and females separately, were then rank-based inverse normalised as one group.

*Number of children (N. children)*

The N. children variable was constructed from answers to the touchscreen questions ‘How many children have you fathered?’ (data-field 2405: ‘Number of children fathered’) and ‘How many children have you given birth to? (Please include live births only)’ (data-field 2734: ‘Number of live births’). Both questions were asked in the initial assessment visit. Data-field 2405 was only collected from males and data-field 2734 was only collected from women. In the subset of 271,317 WB ‘unrelated’ individuals with applicable N. children entries, we adjusted for year of birth up to the order of three, age at recruitment and 40 PCs separately for each sex by performing linear regression in R. The resulting residuals, attained for males and females separately, were then rank-based inverse normalised as one group.

*Number of siblings (N. siblings)*

The N. siblings variable was constructed from answers to the touchscreen questions ‘How many sisters do you have? (Please include those who have died, and twin sisters. Do not include half-sisters, step-sisters or adopted sisters)' (data-field 1883: 'Number of full sisters') and 'How many brothers do you have? (Please include those who have died, and twin brothers. Do not include half-brothers, step-brothers or adopted brothers)' (data-field 1873: 'Number of full brothers'). Both questions were asked in the initial assessment visit. In the subset of 268,191 WB 'unrelated' individuals with applicable N. siblings entries, we adjusted for year of birth, age at recruitment up to the order of three and 40 PCs separately for each sex by performing linear regression in R. The resulting residuals, attained for males and females separately, were then rank-based inverse normalised as one group.

*Vitamin D*

The vitamin D variable was based on data-field 30890-0-0: 'Vitamin D', a biochemistry marker measured in blood samples collected at recruitment. In the group of WB 'unrelated' individuals, 24,116 individuals had missing vitamin D values. As indicated by data-field 30895: 'Vitamin D missing reason' and data-field 30896 'Vitamin D reportability', 784 of those were missing because the values were below reportable range and 2 were missing because the values were above the reportable range. We imputed the 'below reportable range' entries to the minimum vitamin D measurement minus 0.001, that is to 9.999 and the 'above reportable range' entries to the maximum vitamin D measurement plus 0.001, that is to 340.001. In the group of 249,079 WB 'unrelated' individuals with applicable values, we adjusted vitamin D levels for year of birth, age at recruitment up to the order of three and 40 PCs separately for each sex by performing linear regression in R treating rank-based inverse normalised vitamin D values (including the imputed ones) as a dependent variable. The resulting residuals, attained for males and females separately, were then rank-based inverse normalised as one group.

*Glucose*

The glucose variable was based on data-field 30740-0-0: ‘Glucose’, a biochemistry marker measured in blood samples collected at recruitment. In the group of WB ‘unrelated’ individuals, 34,783 individuals had missing glucose values. As indicated by data-field 30745: ‘Glucose missing reason’ and data-field 30746 ‘Glucose reportability’, three of those were missing because the values were below reportable range. We imputed the ‘below reportable range’ entries to the minimum glucose measurement minus 0.001, that is to 1.004. In the group of 237,629 WB ‘unrelated’ individuals with applicable values, we adjusted glucose levels for year of birth, age at recruitment up to the order of three and 40 PCs separately for each sex by performing linear regression in R treating rank-based inverse normalised glucose values (including the imputed ones) as a dependent variable. The resulting residuals, attained for males and females separately, were then rank-based inverse normalised as one group.

*Grip strength*

The grip strength variable was based on data-field 46 ‘Hand grip strength (left)’ and data-field 47 ‘Hand grip strength (right)’, physical measures attained in the initial assessment visit. For each individual we chose the maximum value (left, right) as their grip strength measurement. For the 270,525 WB ‘unrelated’ individuals with non-missing values, we adjusted grip strength measurements for year of birth, age at recruitment up to the order of three, standing height and 40 PCs separately for each sex by performing linear regression in R treating rank-based inverse normalised grip strength values as a dependent variable. The resulting residuals, attained for males and females separately, were then rank-based inverse normalised as one group.

*Lipoprotein A*

The lipoprotein A variable was based on data-field 30790-0-0: ‘Lipoprotein A’, a biochemistry marker measured in blood samples collected at recruitment. In the group of WB ‘unrelated’ individuals, 65,941 individuals had missing lipoprotein A values. As indicated by data-field 30795: ‘Lipoprotein A missing reason’ and data-field 30796 ‘Lipoprotein A reportability’, 26,943 of those were missing because the values were below reportable range and 18,287 were missing because the values were above the reportable range. We imputed the ‘below reportable range’ entries to the minimum lipoprotein A measurement minus 0.001, that is to 3.799 and the ‘above reportable range’ entries to the maximum lipoprotein A measurement plus 0.001, that is to 189.001. In the group of 251,698 WB ‘unrelated’ individuals with applicable values, we adjusted lipoprotein A levels for year of birth, age at recruitment up to the order of three and 40 PCs separately for each sex by performing linear regression in R treating rank-based inverse normalised lipoprotein A values (including the imputed ones) as a dependent variable. The resulting residuals, attained for males and females separately, were then rank-based inverse normalised as one group.

*Sex hormone binding globulin (SHBG)*

The SHBG variable was based on data-field 30830-0-0: ‘SHBG’, a biochemistry marker measured in blood samples collected at recruitment. In the group of WB ‘unrelated’ individuals, 36,843 individuals had missing SHBG values. As indicated by data-field 30835: ‘SHBG missing reason’ and data-field 30836 ‘SHBG reportability’, 5 of those were missing because the values were below reportable range and 389 were missing because the values were above the reportable range. We imputed the ‘below reportable range’ entries to the minimum SHBG measurement minus 0.001, that is to 0.389 and the ‘above reportable range’ entries to the maximum SHBG measurement plus 0.001, that is to 241.921. In the group of 235,960 WB ‘unrelated’ individuals with applicable values, we adjusted SHBG levels for year of birth, age at recruitment up to the order of three and 40 PCs separately for each sex by performing linear regression in R treating rank-based inverse normalised SHBG values (including the imputed ones) as a dependent variable. The resulting residuals, attained for males and females separately, were then rank-based inverse normalised as one group.

*Testosterone*

The testosterone variable was based on data-field 30850-0-0: ‘Testosterone’, a biochemistry marker measured in blood samples collected at recruitment. In the group of WB ‘unrelated’ individuals, 36,861 individuals had missing testosterone values. As indicated by data-field 30855: ‘Testosterone missing reason’ and data-field 30856: ‘Testosterone reportability’, 22,009 of those were missing because the values were below reportable range and 13 were missing because the values were above the reportable range. We imputed the ‘below reportable range’ entries to the minimum testosterone measurement minus 0.001, that is to 0.349 and the ‘above reportable range’ entries to the maximum testosterone measurement plus 0.001, that is to 54.343. In the group of 257,570 WB ‘unrelated’ individuals with applicable values, we adjusted testosterone levels for year of birth, age at recruitment up to the order of three and 40 PCs separately for each sex by performing linear regression in R treating rank-based inverse normalised testosterone values (including the imputed ones) as a dependent variable. The resulting residuals, attained for males and females separately, were then rank-based inverse normalised as one group.

*Dietary study invitation*

The dietary study invitation variable was based on data-field 11001: ‘Invitation to complete online 24-hour recall dietary questionnaire, acceptance’. The 166,993 WB ‘unrelated’ individuals with the data-field values ‘No response’, ‘Partial’ or ‘Completed’ were defined as cases. That is, they had been invited to participate in the study. The 105,416 WB ‘unrelated’ individuals with non-applicable (NA) values in this field were defined as controls. That is, they had not been invited to the study.

*Dietary study participation*

The dietary study participation variable was based on data-field 11001: 'Invitation to complete online 24-hour recall dietary questionnaire, acceptance'. The 54,124 WB 'unrelated' individuals with the values 'Partial' or 'Completed' in the field were defined as cases. The 112,869 WB 'unrelated' individuals with the data-field value 'No response' were defined as controls.

*Physical activity study invitation*

The physical activity study invitation variable was based on data-field 11005: 'Invitation to physical activity study, acceptance'. The 132,633 WB 'unrelated' individuals with the data-field values 'No response', 'Partial' or 'Completed' were defined as cases. That is, they had been invited to participate in the study. The 139,776 WB 'unrelated' individuals with non-applicable (NA) values in this field were defined as controls. That is, they had not been invited to the study.

*Physical activity study participation*

The physical activity study invitation variable was based on data-field 11005: 'Invitation to physical activity study, acceptance'. The 59,455 WB 'unrelated' individuals with the data-field values 'Partial' or 'Completed' were defined as cases. The 73,178 WB 'unrelated' individuals with the data-field value 'No response' were defined as controls.

**GWAS summary statistics**

To compute LD-score regression intercepts (see below) we required GWAS summary statistics for the corresponding phenotypes. We attained those statistics by performing GWAS with BOLT-LMM[8] (version 2.3) in the group of 272,409 'unrelated' WB individuals in UKBB. For quantitative phenotypes, which had been transformed as described above (see *Phenotypes*), genotype measurement batch (data-field 22000) was added as an additional covariate. For the categorical phenotypes, year of birth, age at recruitment up to the order of three, 40 PCs and genotype measurement batch were added as covariates. For computing the LD score regression intercepts we used the standard linear regression P-values, P_LINREG, from the BOLT output.

**LD score regression intercepts**

When examining the association between the pPGS and a range of phenotypes in the group of 272,409 WB 'unrelated' individuals (Table 1), we accounted for population stratification by dividing the chi-square statistics with the corresponding LD score regression intercept[9]. We computed LD score regression intercepts with the program LDSC[9] (version 1.0.1) using the European Ancestry LD scores computed by the Pan-UKB team[10] (downloaded on April 7th 2021) and our GWAS summary statistics based on the 'unrelated' WB UKBB samples (see *GWAS summary statistics* above). The LD score regression intercepts were computed from the 443,554 common high-quality SNPs (inclusion criteria explained in *Variant-filtering* above) for which we had LD-scores for.

**Relative allele frequencies in different groups of individuals and segments**

Under very broad conditions, if an allele promotes participation, its frequency in the shared segments of participating first degree-relative pairs would be higher than its frequency in the not-shared segments. One obvious question of interest is the quantitative relationship between this frequency difference and the difference of allele frequencies in participants and non-participants. We investigate that through the simulations described below. It is noted that the simulations are based on simplified models and thus cannot capture the real conditions perfectly. However, from other simulations we have performed, the results are quite robust to modest deviations from the model. In particular, we believe they provide a reasonable guide for understanding the UKBB data.

Assume a population of sib-pairs where the sib-pairs are indexed by $i$ and the sibs within a pair are indexed $j = 1, 2$. The probability for sib *ij* to participate is $P(Y_{ij} = 1) = p_{ij}$. We simulated from the logistic regression model

$$logit(p_{ij}) = \alpha + \beta X_{ij}$$

where

$$X_{ij} = w_1 G_{ij} + w_A A_i + w_B B_{ij}.$$

The components $G_{ij}, A_i$ and $B_{ij}$ were assumed to be independent and each had variance 1. $G_{ij}$ was the standardized genotype of a *SNP,* and $A_i$ and $B_{ij}$ were normally distributed variables. Furthermore, we assumed

$$w_1^2 + w_A^2 + w_B^2 = 1$$

so that $var(X_{ij}) = 1$. $B_{i1}$and $B_{i2}$ were assumed to be independent, and the correlation between $X_{i1}$ and $X_{i2}$ was induced by the shared component $A_i$ and the correlation between $G_{i1}$ and $G_{i2}$. In particular,

$$cor^2(X_{i1}, X_{i2}) = w_A^2 + \frac{w_1^2}{2} .$$

As seen below, both $A_i$ and $B_{ij}$ could incorporate genetic as well as environmental factors. The marginal participation rate $E(p_{ij})$ is a function of $\alpha$ and $\beta$. For the simulations, we chose $\alpha = -3.48$ and $\beta = 1.25$. This ensured that $E(p_{ij}) \approx 0.055$. Note that $\beta$ needs to be big enough to incorporate substantial correlation between $Y_{i1}$ and $Y_{i2}$. The reasons for choosing these specific values are given below. We picked $w_1 = 0.1$ ($w_1^2 = 0.01$) so that the single SNP, $G_{ij}$, accounted for 1% of the variance of $X_{ij}$. With $w_1 = 0.1$, $w_A^2 + w_B^2 = 0.99$. Note that the higher the value of $w_A^2$, the higher is $E(p_{i2}|Y_{i1} = 1)$, the expected conditional probability that sib 2 is a participant given that sib 1 is a participant. One of our simulations was performed based on having

$$w_A^2 = \frac{2}{3} - 0.005, w_B^2 = \frac{1}{3} - 0.005.$$

In combination with $\alpha = -3.48$ and $\beta = 1.25$, the simulations showed that

$$E(p_{i2}|Y_{i1} = 1) \approx 0.110 = 2 \times 0.055,$$

or approximately twice as many sib-pairs were in the sample relative to what was expected from random participation, as in the case of the UKBB samples[1]. While these parameter values can correspond to many scenarios, we had particularly the following in mind. Consider a further decomposition

$$w_A^2 A_i = w_{AI}^2 A_{iI} + w_{AII}^2 A_{iII}$$

where both $A_{i1}$ and $A_{i2}$ are standard normal, and

$$w_{AI}^2 = w_B^2 = \frac{1}{3} - 0.005, w_{AII}^2 = \frac{1}{3}.$$

This corresponds to a scenario where one-third of the variance of $X_{ij}$ is accounted for by a shared environmental (non-genetic) component $A_{iII}$, while $G_{ij}$, $A_{iI}$, and $B_{ij}$ are all genetic. In particular, $A_{iI}$ plus $B_{ij}$ is the genetic component to participation other than $G_{ij}$. On average, one half of that genetic component is shared between the two sibs ($A_{iI}$), and one half is not shared ($B_{ij}$). Note that in this scenario, shared genetics and shared environment contribute equally to the participation correlation of the two sibs.

Let $f$ be the population frequency of the positively selected allele of $G_{ij}$. We used $\Delta_f$ to denote the change of frequency of this allele relative to the population frequency. From the simulations, we calculated $\Delta_f$ for the sample as a whole ($\Delta_f(sample) = f(sample) - f$), for the shared alleles in the participating sib-pairs ($\Delta_f(shared)$), for the not-shared alleles

($\Delta_f(notshared)$), for the participating sib-pairs ($\Delta_f(sibs)$), and for the singletons ($\Delta_f(singletons)$). Of interest were the values of the various $\Delta_f$'s relative to $\Delta_f(sample)$. In the simulation based on the parameter values mentioned above and with $f = 0.5$,

$$\frac{\Delta_f(shared)}{\Delta_f(sample)} = 1.80, \frac{\Delta_f(notshared)}{\Delta_f(sample)} = 0.90, \frac{\Delta_f(sibs)}{\Delta_f(sample)} = 1.35, \frac{\Delta_f(singletons)}{\Delta_f(sample)} = 0.96.$$

Also,

$$\frac{f(shared) - f(notshared)}{f(sample) - f(population)} = \frac{\Delta_f(shared) - \Delta_f(notshared)}{\Delta_f(sample)} = 0.90.$$

As noted above, for the chosen parameter values,

$$\frac{E(p_{i2}|Y_{i1} = 1)}{E(p_{i2})} \approx 2.$$

To understand better how the relative frequency changes are related to the induced enrichment of sib-pairs, we performed simulations with other values of $w_A^2$. In particular, for $\pi = 0.0, 0.1, 0.2, 0.3, 0.4, 0.5, 0.6, 0.7, 0.8, 0.9, 1.0, 1.1, 1.2, 1.3, 1.4,$ and 1.496, we performed simulations with

$$w_A^2 = \pi\left(\frac{2}{3} - 0.005\right).$$

For each value of $w_A^2$, we simulated 500 replications from a population of $10^8$ sib-pairs. In Fig. S9a, we plotted simulated values of

$$\frac{\Delta_f(shared)}{\Delta_f(sample)}, \quad \frac{\Delta_f(notshared)}{\Delta_f(sample)}, \frac{\Delta_f(sibs)}{\Delta_f(sample)}, \frac{\Delta_f(singletons)}{\Delta_f(sample)}$$

as functions of the simulated value of

$$\frac{E(p_{i2}|Y_{i1} = 1)}{E(pi2)}.$$

Note when $w_A^2 = 0$, where $G_{ij}$ accounts for all the correlation between $Y_{i1}$and $Y_{i2}$, $E(p_{i2}|Y_{i1} = 1)$is only a little higher than $E(p_{i2})$, while $\Delta_f(shared)$ is close to twice the value of $\Delta_f(sample)$, and $\Delta_f(notshared)$ is very close to $\Delta_f(sample)$. In Fig. S9b, simulated values of

$$\frac{f(shared) - f(notshared)}{f(sample) - f(population)} = \frac{\Delta_f(shared) - \Delta_f(notshared)}{\Delta_f(sample)}$$

are plotted against simulated values of

$$\frac{E(p_{i2}|Y_{i1} = 1)}{E(pi2)}.$$

**Significant associations in the MHC region**

In our participation GWAS, we excluded variants in the MHC region (chr6:25,000,000-33,500,000) as well as other extended LD regions (Table S2). If we had not applied that criteria, 3,975 variants in the MHC would have been included in our participation GWAS as they passed other variant filters (see *Variant filtering* above). Of those, 204 SNPs associated with ascertainment with $P < 5.0\times10^{-8}$. Further investigations led us to believe that these associations were most likely due to data artefacts in the MHC. Firstly, the significant results were solely driven by the sibling pairs. That is, for these 204 variants, differences between the shared and not shared allele frequencies were only observed among the sibling pairs and not among the parent-offspring pairs (Fig. S6). Secondly, for an ascertainment effect of this degree, we would expect to observe allele frequencies differences between the siblings and the 'unrelated' group. However, no such differences were observed for these variants. Thirdly, the MHC region was as prone to deviations from expected IBD-fractions of sibling pairs as the beginning and ends of chromosomes (Fig. S3) indicating data artefact problems in the area.

**The effects of selection on the associations between trait and genotypes**

Suppose there is biased sample selection with respect to a trait $Y$, directly or indirectly through a correlated trait. The type of biased selection studied here usually would decrease the sample variance of $Y$. If $Y$ is correlated with a genotype $g$ in the population, it is rather common knowledge that $r^2$ between $Y$ and $g$ would be reduced in the sample. What is not quite common knowledge is that biased sample selection can create artificial epistatic effects for variants that have an effect on the trait. For example, suppose the relationship between trait $Y$ and the genotypes of two variants, $g_1$ and $g_2$, is additive in the population:

$$Y = a_1 g_1 + a_2 g_2 + noise.$$

Suppose, $a_1$ and $a_2$ are both positive. In the biased sample, not only would the expected values of the fitted coefficients for $g_1$ and $g_2$ shrink, the expected value of the fitted coefficient for the interaction term $g_1 g_2$ would be non-zero and positive. This can be easily confirmed by simulations.
